# Dynamics of feeding behaviour and meal patterning in protein-restricted mice

**DOI:** 10.64898/2026.05.07.723245

**Authors:** Hamid Taghipourbibalan, James Edgar McCutcheon

## Abstract

Of the three dietary macronutrients, protein plays an especially pivotal role in physiological functions. Nevertheless, the behavioural control of protein intake is poorly understood. In this study, we used Feeding Experimentation Devices (FED3s) to examine the structure of ingestive behaviour in mice given access to diets varying in protein content. Adult C57BL/6NRj mice were contact-housed in pairs in custom-made cages with perforated dividers, each having access to an individual FED3 unit. Mice were given ad libitum access to either 20 mg control, non-restricted (NR) pellets (20% casein) or 20 mg protein-restricted (PR) pellets (5% casein) from FED3s on free-feeding mode. Each pellet retrieval event was timestamped ∼24 h/day. All mice experienced both diets for 7 days with order of diet presentation counterbalanced (i.e., NR→PR and PR→NR). Analysis of dynamics of pellet intake per day revealed that mice that were initially protein-restricted first showed a decrease in pellet intake before increasing on later days and exhibiting a persistent high level of intake once non-restricted diet was available. The group that was initially non-restricted exhibited a blunted response to the same diet manipulation. In addition, we clustered pellet retrieval data into discrete clusters of feeding events and used a mathematical approach to determine the boundary of meals (2-5 pellets), separated from “snacks” (1 pellet) and “feasts” (>5 pellets). We identified alterations in meal patterning in response to diet manipulation with protein restriction increasing “snacking” and leading to increased meal number, and reduced meal size. Moreover, restored access to NR diet, elicited “feasting”. These effects depended on the sequence of diets the mice experienced, such that the effects were stronger in initially protein restricted mice compared to those initially non-restricted. In summary, our findings show that manipulation of dietary protein levels affects meal patterning in adult mice.

## 1. Introduction

Animals alter food intake in response to their physiological state, and it is proposed that behaviour is often targeted towards a nutrient that may be lacking (Watts et al., 2022). Evidence from a range of species suggests that, under many conditions, food-directed behaviour is directed towards achieving a specific protein intake target (Simpson & Raubenheimer, 2005; Sørensen et al., 2008). Consistent with this idea, studies show that growing animals consume greater amounts of protein to support tissue development (Musten et al., 1974), and that protein consumption increases during pregnancy and lactation (Cohen & Woodside, 1989). Conversely, insufficient protein intake can have significant physiological consequences. For example, maternal protein restriction has long-term effects on offspring, predisposing them to increased body weight and metabolic dysfunction in adulthood, including glucose intolerance and hyperinsulinaemia, and is associated with alterations in regulatory pathways such as leptin signalling (Qasem et al., 2016; Zheng et al., 2021).

Dietary protein restriction triggers adaptations in food intake, metabolism, and circulating hormones, which can lead to changes in protein preference and balance of energy expenditure (Morrison & Laeger, 2015). These effects are distinct from caloric restriction alone, for example, protein restriction – but not caloric restriction – increases fibroblast growth factor 21 (FGF21), a hormone that acts centrally to induce protein appetite. In fact, disruption of FGF21 signalling, including loss of FGF21 or its brain co-receptor β-Klotho, blunts the normal adaptive increase in protein-directed intake during protein restriction (Hill et al., 2019). In addition, restriction of essential amino acids affects various metabolic parameters such as feed efficiency, energy expenditure, and glucose metabolism in mice (Yap et al., 2020). Taken together, these findings suggest a sensitive thresholding system to counteract divergence in protein status from its required range.

Understanding how the drive to acquire protein manifests in behavioural dynamics of food intake is important for understanding this form of nutritional regulation. We and others have demonstrated that protein-restricted rats and mice increase their food intake and show a strong preference for protein compared to carbohydrate (Hill et al., 2019; Murphy et al., 2018; Volcko & McCutcheon, 2022). All existing studies on protein appetite, to our knowledge, have only evaluated food intake by measuring the amount of food taken over a period of time, e.g., grams per day. However, this approach does not generate data with high temporal resolution, which is necessary to study aspects of feeding behaviour such as meal patterning.

Here, we use the third version of Feeding Experimentation Devices, FED3, (hereafter FED; Matikainen-Ankney et al., 2021) to accurately monitor feeding in the home-cage allowing analysis of detailed patterns of feeding behaviour in response to protein restriction. We analysed food intake in mice under varying levels of dietary protein and found that protein-restricted mice exhibit an altered feeding structure that is dependent on the sequence of diets they experience. By analysing meal patterning, we divided eating events into fundamental units (e.g., snacks, meals and feasts) and found that behavioural alterations in response to protein restriction are reflected in these elements. These changes to eating patterns may contribute to the compensatory increased food intake or food preferences observed in protein-restricted animals and will help understand the underlying regulatory mechanisms of protein balance.

## 2. Materials and methods

### 2.1. Animals

Adult male and female C57BL/6NRj mice (6-8 weeks old, weighing 20-30 g) purchased from Janvier Labs (France), acclimatized for 5 days in a temperature (22 ± 0.5 °C) and humidity (56 ± 2%) controlled room. The light:dark cycle was maintained on a 12:12 basis (lights on at 07:00 am). Subsequently, mice were moved to a colony room with similar environmental conditions where the experiments were conducted. Water was provided ad libitum at all times. Food was available ad libitum throughout experiments but mode of delivery switched from standard food hoppers to FEDs when experiments started (see below for details). The current study comprised one cohort of male mice (n=12) and one cohort of female mice (n=11) each carried out at relatively close time points (∼3 weeks apart). All animal care and experimentation were in compliance with the EU directive 2010/63/EU for animal experiments and were approved by the Norwegian Food Safety Authority (FOTS protocol #22315).

### 2.2. Housing and home-cage monitoring of ingestive behaviour

Two animals per cage were contact-housed in modified conventional cages (1284L EUROSTANDARD TYPE II L, TECNIPLAST, Italy; 365 x 207 x 140 mm; floor area of 530 cm^2^) separated by a perforated divider that allowed visual, olfactory and minor physical communication but no major physical contact. Each cage was equipped with two FEDs mounted on the outer sides of the cage such that each mouse had access to its own FED unit (Fig. 1A and 1B). FEDs are automated food dispensers that release 20 mg food pellets and monitor feeding behaviour by accurately timestamping every pellet delivery event or other operant behaviours and interactions made to acquire a food reward such as number of nose-pokes or duration of pellet retrieval (Matikainen-Ankney et al., 2021). Almost no intervention by the experimenter is needed to feed the animal and the battery-powered device can run 24h/day, 7 days/week for as long as desired.

**Fig. 1.**
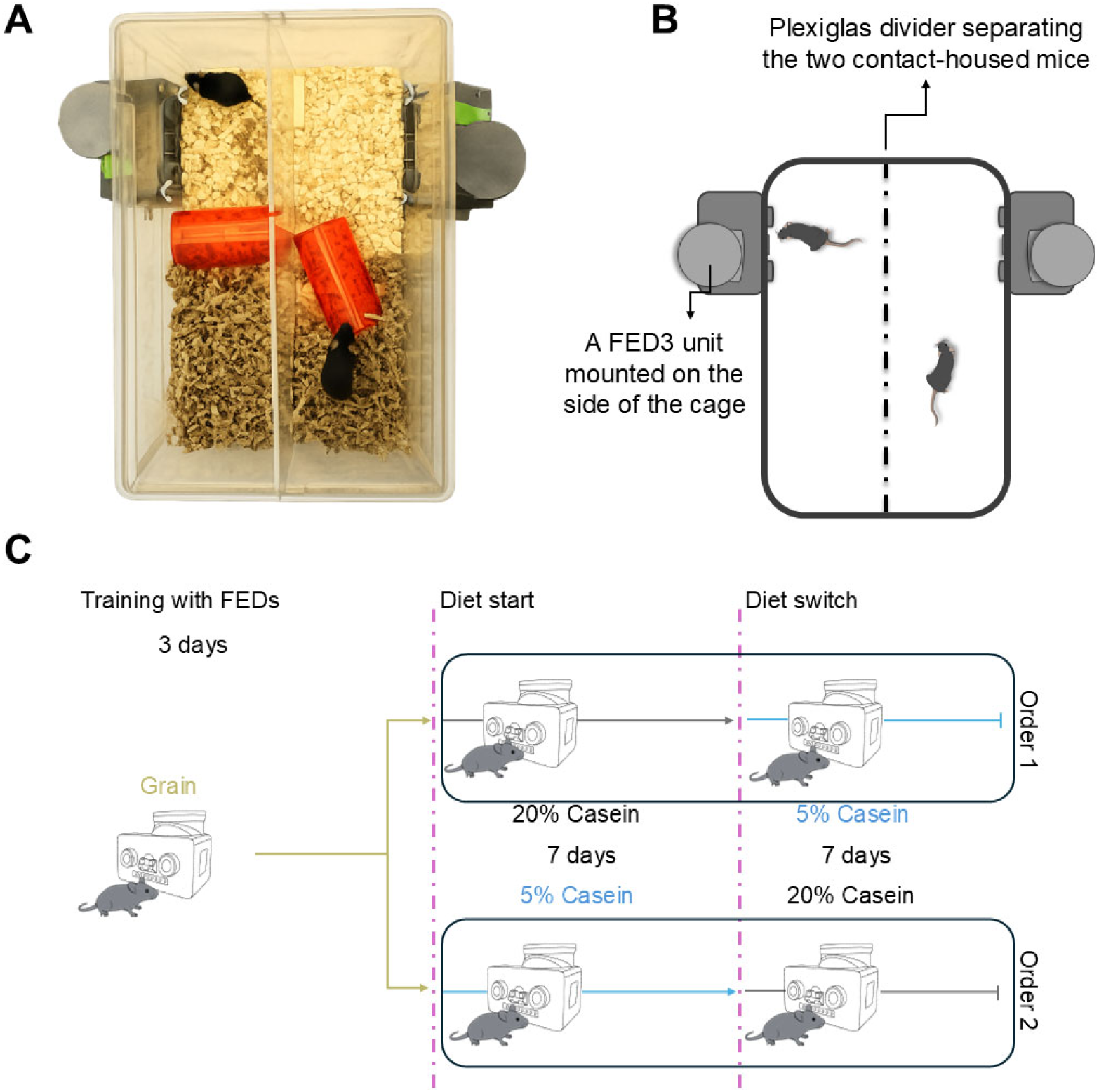
(A) An image of a FED cage with two contact-housed mice, nesting material and enrichment tubes. (B) The diagram of a FED cage displaying the positioning of FED units on the side of the cage and the perforated Plexiglas divider. (C) Schematic representation of the study paradigm. All mice habituated to FEDs during a 3-day training phase where they learned to receive 20 mg grain pellets from the device. After training, mice were randomly grouped to receive either 20% casein pellets (NR) or 5% casein pellets (PR). After one week, diets were switched (NR→ PR, Order 1 and PR→NR, Order 2) and mice were maintained on the new diet for one more week.

### 2.3. Diet manipulation

First, mice were habituated to FEDs for 3 days during which time they learned to collect 20 mg grain pellets (#F0163; Bio-Serv; 3.35 kcal per pellet) from the devices. On the first day of habituation, mice also had access to regular chow pellets in the standard food hopper. After habituation, mice were randomly grouped in equal numbers to either receive non-restricted control diet (NR: nutritionally-balanced pellets with 20% casein, #F07736; Bio-Serv; 4.01 kcal per pellet) or isocaloric protein-restricted diet (PR: similar to NR diet but with 5% casein #F07737; Bio-Serv; 3.99 kcal per pellet) from FEDs (Table S1. Diet formulation). Fat was held at the same level in both pellets (10%), so PR pellets contained additional carbohydrate to replace the missing protein. After one week, diets were switched for all mice such that previously non-restricted mice received protein-restricted pellets and vice versa (Order 1: NR→ PR; Order 2: PR→NR). Hence, all mice experienced each diet for 7 days (14 days in total) with FEDs set on free-feeding mode, i.e., once a pellet was taken, a new pellet was automatically dispensed, and every pellet delivery event was logged ∼ 24 h/day (Fig. 1C). FEDs were maintained every day between 08:00-09:00 at the same time as bodyweight measurement to ensure enough pellets were available and the devices were working reliably.

### 2.4. Data analysis and statistical approaches

Data from the FED3 devices were collected as .csv files and processed in Python (v3.12.2) using custom analysis scripts to extract inter-pellet intervals, compute pellet intake per unit time, and quantify meal pattern parameters. All statistical analyses were conducted in R (v4.4.1) on a 64-bit Windows platform.

Different statistical approaches were applied depending on the structure of each dataset. For Diet-averaged measures (i.e., across NR and PR), including total pellets consumed and the number, size, and frequency of feeding events (snacks, meals, feasts), as well as bodyweight, we used two- or three-way ANOVAs. Order (NR→PR [Order 1] vs PR→NR [Order 2]) and Sex (Male vs Female, when not pooled) were between-subjects factors, and Diet (NR vs PR) was a within-subject factor. Where significant main effects or interactions were detected, post hoc comparisons were performed with Holm’s correction for multiple testing.

To assess temporal dynamics across days, we used mixed-design repeated-measures ANOVA with Diet (NR, PR) and Day (0-6) as within-subject factors, and Order and, where relevant, Sex as between-subjects factors. All effects that included Day used Greenhouse-Geisser corrections; we report the corrected degrees of freedom with the corresponding *F* and *p* values for between-subject effects and two-level within-subject factors (Diet) are reported with uncorrected df. Planned and exploratory simple-effects were conducted with *emmeans* in three families: (1) NR vs PR at each Day (within Order), (2) day-to-day tests within each Diet (both consecutive and all-pairs), and (3) Order-wise differences within each Diet×Day cell. *P*-values were Holm-adjusted within each family. Effect sizes are reported as partial η^2^ (ηp^2^).

For hourly (circadian) outcomes (e.g., pellets per hour), we fitted Linear Mixed-Effects Models with fixed effects of Hour, Diet, Order, and Sex (when relevant), and a random intercept for Mouse to account for repeated measures. Estimated marginal means (EMMs) were computed for targeted contrasts across Hour or Diet, with Holm-adjusted post hoc tests for pairwise comparisons.

A comprehensive list of R packages, statistical functions, and analytical workflows used is provided in Supplementary section 2.4.S1. All data and custom analysis scripts are available at Github (https://github.com/jaimemcc/FEDProtein_dynamics_paper/tree/dynamics_paper).

## 3. Results

### 3.1. Protein restriction dynamically alters food consumption

We first examined whether type of Diet and Order of presentation affected total pellet intake (Fig. 2A). Repeated-measures ANOVA showed a significant main effect of Order (F_1,42_=5.70, p=0.021), indicating that mice in Order 2 consumed more pellets overall, compared to those in Order 1. However, total pellet intake summed across each phase was not significantly affected by Diet (F_1,42_=0.86, p=0.36), and there was no significant interaction between Order and Diet (F_1,42_=1.88, p=0.18).

**Fig. 2.**
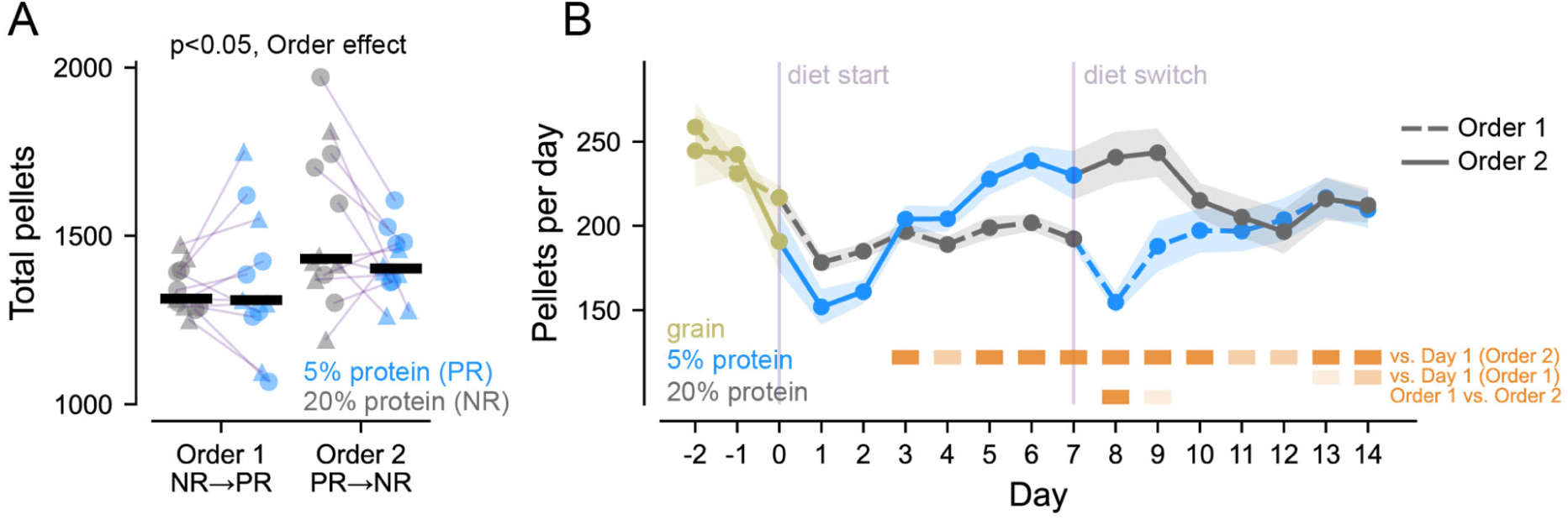
Protein restriction affects food intake in an order-dependent manner. **(A)** Mice that experienced protein-restricted (PR) diet before non-restricted (NR) diet (Order 2) took more food pellets, relative to Order 1 mice. Markers show individual mice with circles as males and triangles as females. Thick black line is median. **(B)** Day-to-day analysis shows that there were diet-induced changes not apparent after averaging across all days in each phase. Order 2 mice showed an initial decrease in pellets per day that increased towards the end of the PR phase, and this increased intake remained high even when NR diet was presented. Order 1 mice did not exhibit the same behaviour, with relatively steady intake across both phases and a slight increase at the end of their PR phase. Data are mean ± SEM with dashed line as Order 1 and solid line as Order 2. Orange symbols show post hoc results with light orange as p<0.05, mid-orange as p<0.01, and dark orange as p<0.001.

Next, instead of only considering total pellet intake across each diet phase, we examined day-to-day variation in pellet intake (Fig. 2B). Data are pooled across sexes, as no significant sex differences were detected; sex-specific data are shown in Fig. S1. Analysis of these dynamics showed a robust main effect of Day, (F_3.87,81.19_ = 11.40, p < 0.001, ηp^2^ = 0.352), with no main effect of Diet (F_1,21_ = 1.19, p = 0.287, ηp^2^ = 0.054), and a marginal trend for Order (F_1,21_ = 4.17, p = 0.054, ηp^2^= 0.166). In addition, there was a strong Diet × Day interaction, (F_2.58,54.23_ = 10.94, p < 0.001, ηp^2^ = 0.343), and a significant three-way Order × Diet × Day interaction, (F_2.58,54.23_ = 4.91, p = 0.006, ηp^2^ = 0.190). Other interactions were not significant (Order × Day: F_3.87,81.19_ = 0.87, p = 0.485, ηp^2^ = 0.040; Order × Diet: F_1,21_ = 2.98, p = 0.099, ηp^2^ = 0.124).

Post hoc tests showed that the significant Diet × Day and Order × Diet × Day interactions are driven almost entirely by Order 2 mice (PR→NR). In Order 2, there appears to be an immediate suppression of pellet intake (likely, linked to changing caloric value of the pellets, see Fig. S2) but after two days, pellet intake rises steadily by mid/late PR and remains high. In fact, after switching back to NR, pellet intake remains high and stays elevated, relative to Day 1, for several days before declining. This pattern, particularly the gradual increase while restricted, likely reflects adaptation to low-protein diet via hyperphagia and the sustained elevation of intake after switching back to NR diet is compensatory feeding to replenish protein. Surprisingly, in Order 1 mice, except for the initial dip after switching to PR diet, this hyperphagia was not observed. In summary, pellet consumption varied across days, and the day-wise pattern differed by Diet. Moreover, the Diet-dependent time course itself varied by the order of diet presentation.

### 3.2. Sex dependent effects of protein restriction on bodyweight

Bodyweight trajectories across diet phases were analysed using repeated-measures ANOVA with Sex, Order, and Diet as between-subject factors and Day as a within-subject factor (Fig. 3A and 3B). There was a strong main effect of Sex (F_1,19_ = 75.69, p < 0.001, ηp^2^ = 0.799), indicating marked overall differences in bodyweight between male and female mice. A significant main effect of Diet was also observed (F_1,19_ = 4.68, p = 0.043, ηp^2^ = 0.198), showing that bodyweight differed between PR and NR conditions. In addition, bodyweight changed significantly across days (F_2.91,55.30_ = 3.79, p = 0.016, ηp^2^ = 0.166), while the main effect of Order did not reach significance (F_1,19_ = 3.19, p = 0.090, ηp^2^ = 0.144).

**Fig. 3.**
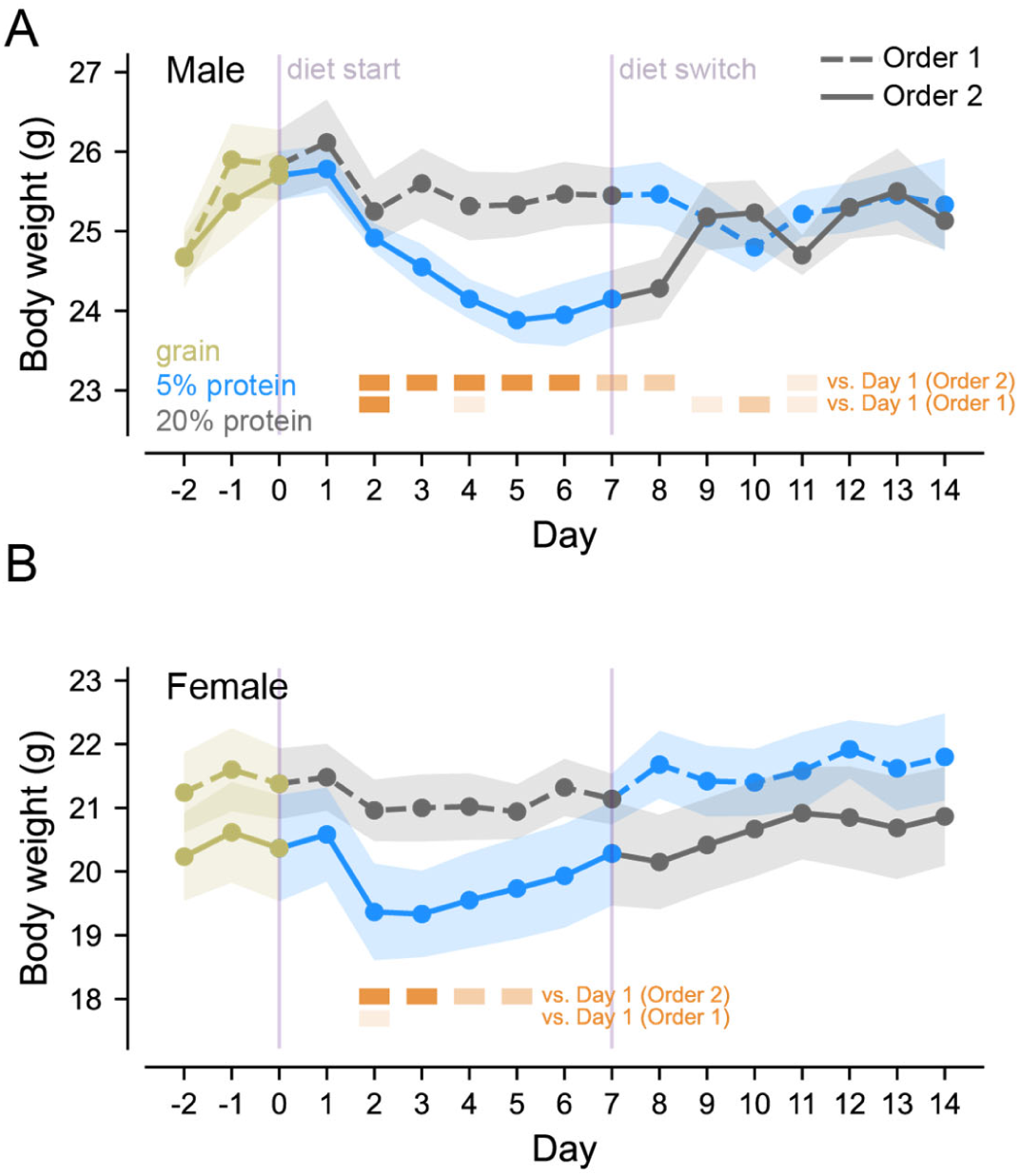
Bodyweight differed between PR and NR conditions in a sex-dependent manner. **(A)** In male mice, those that received PR diet first (Order 2), experienced a marked drop in bodyweight during the PR phase followed by weight regain in the NR phase, but this effect was very mild in Order 1 male mice. **(B)** In female mice, PR diet caused a moderate and transient reduction in body weight in Order 2 mice, followed by bodyweight regain even before switching to NR diet. Bodyweight of Order 1 female mice remained almost resistant to diet manipulation. Statistical annotation as in Fig. 2B.

Importantly, Diet interacted with Order (F_1,19_ = 9.73, p = 0.006, ηp^2^ = 0.339), indicating that the effect of diet on bodyweight depended on the sequence in which diets were experienced. A significant Diet × Day interaction was also detected (F_2.08,39.59_ = 4.90, p = 0.012, ηp^2^ = 0.205), showing that bodyweight trajectories changed over time differently across diet conditions. Moreover, a robust Order × Diet × Day interaction was found (F_2.08,39.59_ = 10.80, p < 0.001, ηp^2^ = 0.362), demonstrating that the day-by-day pattern of bodyweight change varied as a function of diet-switch order. No other interactions were significant (all p ≥ 0.10), although there were trends for Sex × Day (F_2.91,55.30_ = 2.51, p = 0.070, ηp^2^ = 0.117) and Order × Sex × Diet (F_1,19_ = 3.81, p = 0.066, ηp^2^ = 0.167).

Post hoc comparisons showed that male mice in Order 2 exhibited the clearest bodyweight reduction during the PR phase, followed by recovery after switching to the NR diet. Female mice in Order 2 displayed a milder decrease in bodyweight and a faster recovery. In addition, male mice in Order 1 also showed some reduction in bodyweight after transitioning to the PR phase, although this effect was less pronounced than in Order 2 males. Together, these findings indicate that bodyweight dynamics were shaped not only by diet condition, but also by the order of dietary exposure, with male mice showing the strongest sensitivity to protein restriction.

### 3.3. Protein restriction does not affect general circadian rhythms

Next, we used a mixed-effects model to explore the circadian pattern of food intake across each diet phase (Fig. 4). As expected, analysis showed a main effect of Hour (F_23,893_ = 200.69, p <0.001) highlighting a circadian pattern with increased food intake in the first half of the dark phase and the highest peaks during Hour 14, 15, and 16 (2-4 h after lights out). No significant difference between male and female mice was observed (F _1,19_ = 0.75, p = 0.39). There were trends towards a main effect of Diet (F_1,893_ = 3.50, p = 0.062) and Order (F_1,19_ = 4.29, p = 0.052) and a significant interaction between Diet × Order (F_1,19_ = 9.46, p = 0.002). Post hoc comparisons did not show significant pairwise differences suggesting that the interaction is driven by overall patterns rather than sharp contrasts at specific levels. Moreover, an interaction between Order × Hour (F_1,893_ = 3.56, p <0.001) was identified and post hoc tests showed that Order 2 mice in particular increased intake at peak hours of the NR phase (Fig. 4).

**Fig. 4.**
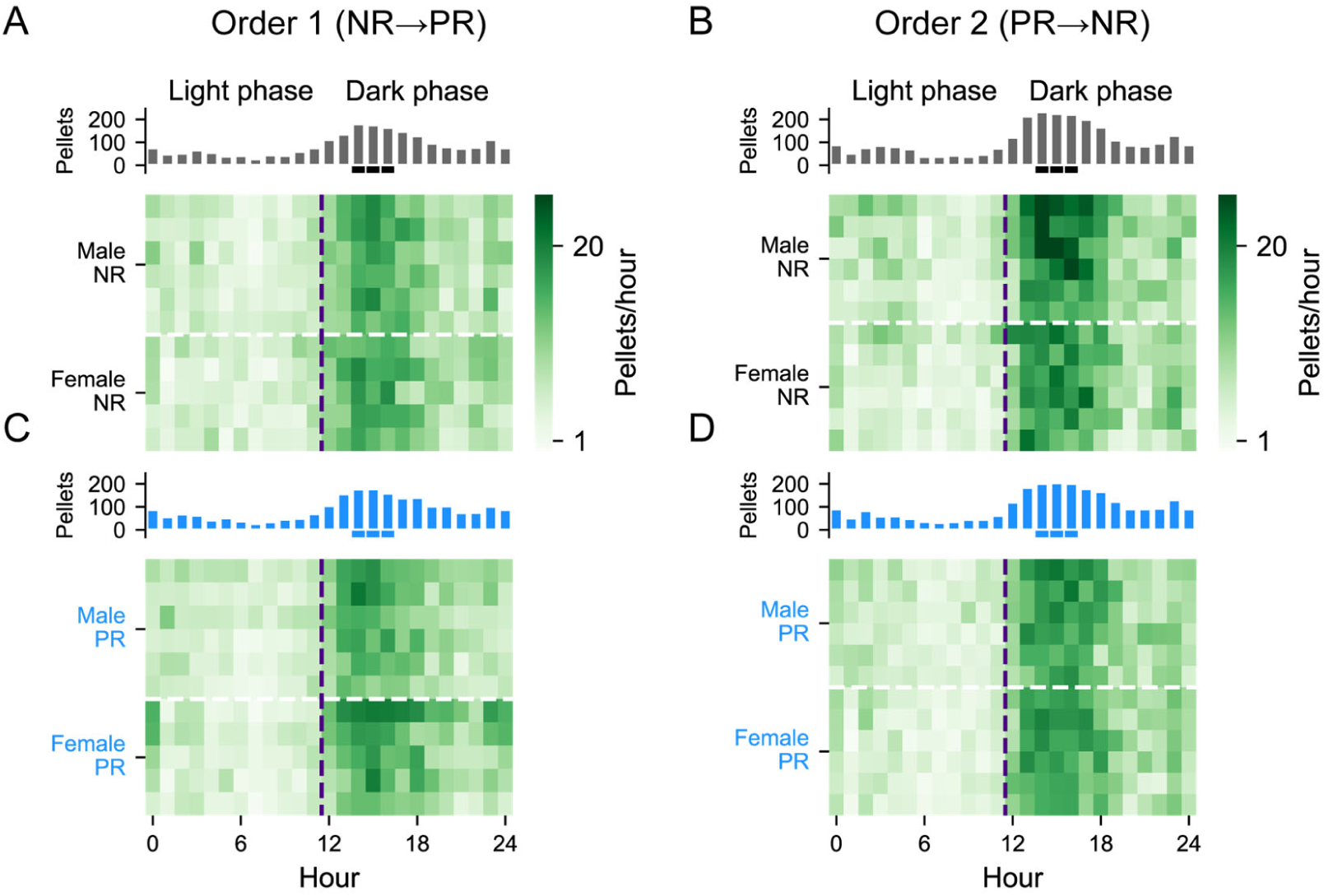
Circadian patterns of food intake show an expected peak of nocturnal feeding. **(A)** Average daily food intake for male and female mice in Order 1 (NR→PR) while on non-restricted (NR) diet. Top panel, bars show summed pellet intake with peak hours highlighted by thick lines underneath bars. Lower panel, heat map where rows are individual male mice (upper rows) and female mice (lower rows). **(B)** As in A but for mice in Order 2 (PR→NR). **(C, D)** As in A and B but for male and female mice on protein-restricted diet.

These findings align with the expected nocturnal feeding behaviour of mice, however, Order 2 showed higher overall pellet intake compared to Order 1, particularly when on the NR diet. Hourly pellet intake did not differ significantly between males and females, and no significant Diet × Sex interaction was detected. These findings suggest that with free access to food on the free feeding mode of FED devices, protein restriction does not affect the circadian rhythm of feeding in mice. Rather, elevated feeding takes place during the regular peak hours of food intake instead of spreading across the day.

### 3.4. Definition of a meal and analysis of feeding microstructure

Each FED3 unit records feeding-related interactions, including pellet retrievals and nose pokes. To define feeding bouts, we analysed the distribution of inter-pellet intervals calculated from consecutive pellet retrieval timestamps and used this distribution to establish a threshold for grouping pellets into discrete feeding events. This analysis focused on the NR phase of Order 1 mice, with males and females combined, as this condition served as a reference group in our experiment. Males and females were pooled as we did not find differences in pellet intake (Fig. 2) although analysis divided by sex is included in Supplementary Material. We used the density estimation of all inter-pellet intervals (IPIs) to divide pellet retrievals into eating events. Our IPI data are in broad agreement with earlier FED studies (Barrett et al., 2025; Matikainen-Ankney et al., 2021), and show that the majority of IPIs were <60 s (Fig. S3). Thus, an *IPI* > 1 min represents a temporal threshold that can be used to determine the end of an eating event.

Using this 60 s criterion, we grouped pellet retrievals into completed feeding clusters for each mouse and examined the final size of each cluster, expressed as the number of pellets retrieved within that cluster. The completed cluster-size distribution showed that 1-pellet clusters were common, 2 to 5-pellet clusters formed the principal multi-pellet range, and clusters larger than 5 pellets were less frequent extended clusters (Fig. S4A; Table S2).

The decline in completed cluster frequency with increasing cluster size was relatively well described by an exponential decay function, indicating that larger feeding clusters became progressively less frequent (Fig. S4A). However, some cluster sizes did not follow the predicted exponential pattern as closely as others. Therefore, to identify demarcations between different cluster-size ranges, we calculated neighbouring completed cluster-size ratios. Neighbouring cluster-size ratios described how frequent completed clusters of size *x*+ 1 were relative to clusters of size *x*. This analysis highlighted the 1 to 2 and 5 to 6 transitions, corresponding to the separation between isolated single pellet events and multi-pellet events, and between the 2-5 pellet range and the >5 pellet extended-cluster range, respectively (Fig. S4B; Table S3).

We also calculated cluster-continuation probability as *P*(*reac*ℎ *x* + 1 l *reac*ℎ*ed x*), asking, among feeding clusters that had reached pellet *x*, what proportion also reached pellet *x* + 1 (Fig. S4C; Table S4). This analysis described stepwise progression across pellet numbers and showed substantial continuation through the 2-5 pellet range. Based on these combined observations, we classified feeding events into three operational categories; “snacks” consisting of a single pellet, “meals” consisting of 2-5 pellets, and “feasts” consisting of more than 5 pellets (Fig. S4D; Table S5).. Using criteria derived from the reference group, we then examined how protein restriction affected each class of feeding event. A detailed explanation of the approach is provided in Supplementary section 3.4.S1.

### 3.5. Distribution of meals (2-5 pellets)

To investigate whether dietary protein restriction alters the patterning of meals, we analysed number of meals and size of meals. Our first analysis considered the average of these parameters across each diet phase, and the second considered day-to-day dynamics within each phase. The analysis of the average number of meals (Fig. 5A) taken over each phase showed a main effect of Diet (F_1,42_ = 10.57, p = 0.002), no main effect of Order (F_1,42_ = 2.71, p = 0.107) and no interaction of Order × Diet (F_1,42_ = 1.52, p = 0.225). Analysis of meal size (Fig. 5C) showed a main effect of Diet (F_1,42_ = 20.55, p < 0.001), no main effect of Order (F_1,42_ = 0.40, p = 0.529), but there was a significant Order × Diet interaction (F_1,42_ = 5.03, p = 0.030). Post hoc tests on meal size revealed a significant decrease in meal size for Order 2 mice when on PR diet, relative to their NR phase (p < 0.001), and relative to Order 1 PR phase (p < 0.01). Thus, mice on PR diet took more frequent meals than when on the NR diet, but these meals were smaller. These effects tended to be greater in Order 2 mice than in Order 1 mice.

**Fig. 5.**
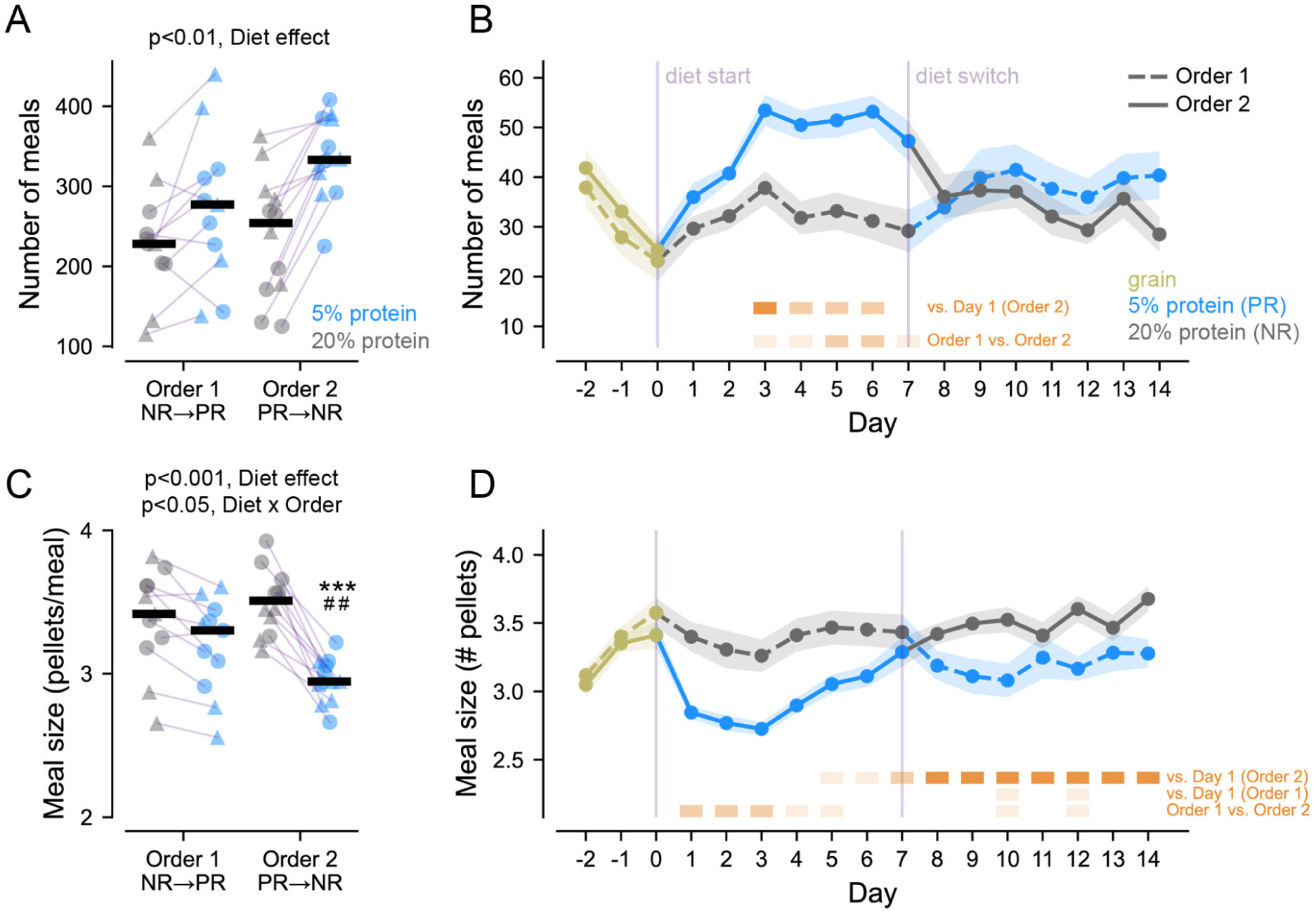
Protein restriction reshapes meal patterning in an order-dependent manner. **(A)** When averaged across phases, mice on the protein-restricted (PR) diet took more meals than when on non-restricted (NR) diet. **(B)** Day-to-day analysis showed that this pattern was most apparent in Order 2 mice, that showed an increase during PR phase and a decrease during the NR phase. **(C)** Meal size was smaller for mice on PR diet than on NR diet and this effect was driven by changes in Order 2 mice. ***, p<0.001 vs. NR phase; ##, p<0.01 vs. Order 1 mice on PR diet. **(D)** Day-to-day analysis showed that this pattern was more apparent in Order 2 mice with a reduction in meal size during the PR phase and an increase during the NR phase. Statistical annotation and plotting conventions follow those described in the caption of Fig. 2.

Next, we examined day-to-day changes in meal number across the two weeks (Fig. 5B). Order 2 mice showed a clear ramp within PR, followed by a drop after switching to NR. By contrast, Order 1 mice varied modestly across days and meal number stayed relatively steady across both phases. A mixed-design repeated-measures ANOVA confirmed main effects of Diet (F_1,21_ = 32.86, p < 0.001, ηp^2^ = 0.610) and Day (F_3.35,70.44_ = 6.28, p < 0.001, ηp^2^ = 0.230), no main effect of Order (F_1,21_ = 1.74, p = 0.201, ηp^2^ = 0.077), and significant Diet × Day (F_4.06,85.30_ = 3.55, p = 0.010, ηp^2^ = 0.145) and Order × Diet interactions (F_1,21_ = 4.58, p = 0.044, ηp^2^ = 0.179). The three-way Order × Diet × Day interaction was also significant (F_4.06,85.30_ = 3.29, p = 0.014, ηp^2^ = 0.135). Holm-corrected post hoc tests showed that within Order 2, meal number increased across early PR days, whereas no day-to-day contrasts survived correction in Order 1.

Examination of day-to-day changes in meal size (Fig. 5D) showed that Order 2 mice tended to have smaller meals during PR and larger meals in the NR phase, whereas Order 1 mice showed more modest day-to-day variation. ANOVA showed a main effect of Diet (F₁,₂₁ = 116.02, p < 0.001, ηp^2^ = 0.847) and a strong Order × Diet interaction (F_1,21_ = 26.51, p < 0.001, ηp^2^ = 0.558), alongside a robust Day effect (F_4.78, 100.40_ = 9.00, p < 0.001, ηp^2^ = 0.300) and an Order × Day interaction (F_4.78, 100.40_ = 2.56, p = 0.034, ηp^2^ = 0.109). The Diet × Day term showed a trend but did not reach significance (F_4.51,94.70_ = 2.03, p = 0.088, ηp^2^ = 0.088), and the three-way interaction was not significant either (F_4.51,94.70_= 1.66, p = 0.160, ηp^2^ = 0.073). See Fig. S5 and S6 for sex-specific plots of Fig. 5B and Fig. 5D)

Together, these results indicate that prior diet exposure (i.e., whether mice started on PR or NR diet) shaped how meal patterning evolved across days, with pronounced adaptations in Order 2 mice during PR and notable shifts after switching to NR diet. Mice that started on PR began with smaller meals early in PR and then steadily increased, with further increases after the switch to NR. By contrast, mice that started on NR showed a similar pattern but with far more modest variation where specific contrasts did not reach significance.

### 3.6. Distribution of “snacks” (1 pellet)

We repeated the analysis used to compare meal patterns above but considering “snacks” (i.e., eating events consisting of just one pellet). When analysing the average number of snacks (Fig. 6A) the two-way ANOVA indicated a main effect of Diet (F_1,42_ = 17.91, p = 0.001), no main effect of Order (F_1,42_ = 2.65, p = 0.111) and an Order × Diet interaction (F_1,42_ = 6.97, p = 0.012). Post hoc comparisons confirmed that Order 2 mice consumed significantly more snacks in the PR phase compared with their NR phase (p < 0.001) and both the PR and NR phases in Order 1 mice (p < 0.05 and p < 0.01, respectively).

**Fig. 6.**
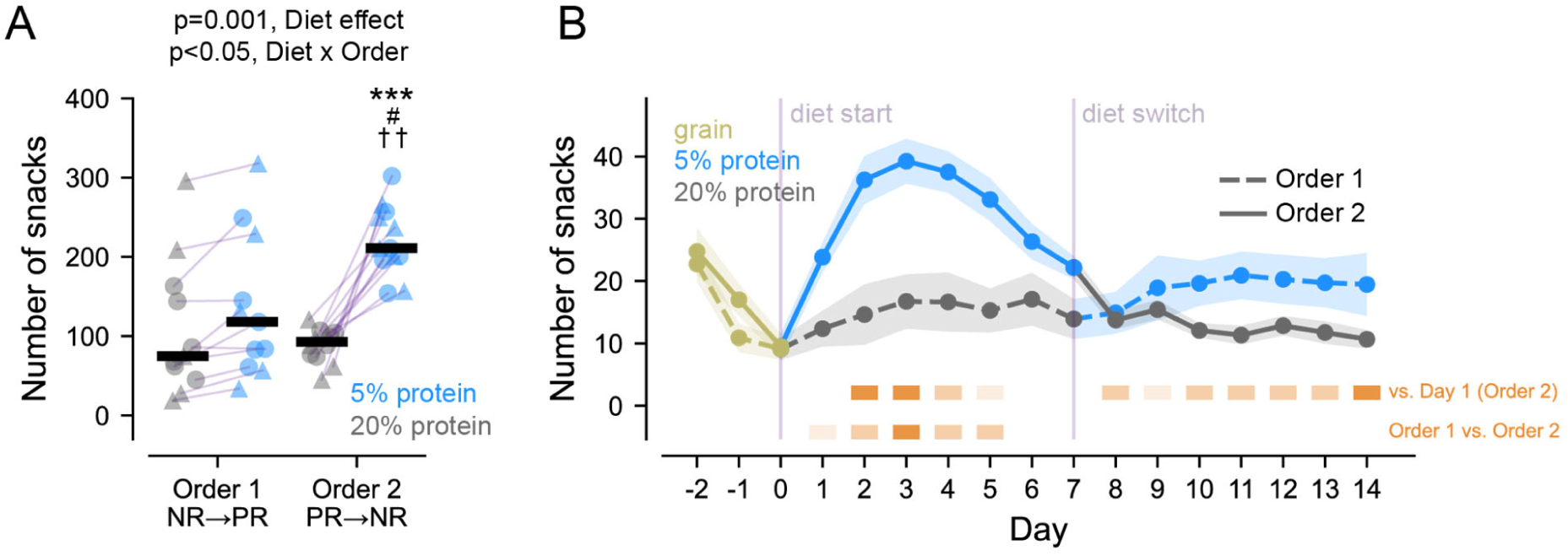
Protein restriction increases snack consumption in an order-dependent manner. **(A)** When averaged across phases, mice consumed more snacks when on the PR diet compared to the NR diet. These effects were primarily driven by Order 2 mice, who displayed a marked rise in number of snacks during the PR phase, whereas Order 1 mice showed only minor changes across diets. Post hoc tests confirmed that Order 2 mice took significantly more snacks during PR than in their NR phase, and compared to both PR and NR phases of Order 1. ***, p<0.001 vs. NR phase; #, p<0.05 vs. Order 1 mice on PR diet; ††, p<0.01 vs. Order 1 mice on NR diet. **(B)** Day-to-day analyses confirmed this pattern with Order 2 mice showing a clear increase in snack number for most days in the PR week, followed by a decrease in snacking after switching to NR. In contrast, Order 1 mice exhibited modest and inconsistent day-to-day variations with no sustained diet-related trend. Statistical annotation and plotting conventions follow those described in the caption of Fig. 2.

Analysis of day-to-day changes in the number of snacks across the two weeks (Fig. 6B), showed that mice that started on PR (Order 2) showed a pronounced rise in snacks during PR with fewer snacks after switching to NR, whereas mice that started on NR varied more modestly across days. ANOVA confirmed a large main effect of Diet (F_1,21_ = 83.09, p < 0.001, ηp^2^ = 0.798), and a strong Order × Diet interaction (F_1,21_ = 35.69, p < 0.001, ηp^2^ = 0.630). There was also a Day effect, (F_3.51,73.67_ = 5.63, p < 0.001, ηp^2^ = 0.211), with significant Order × Day (F_3.51,73.67_ = 2.84, p = 0.036, ηp^2^ = 0.119), and Diet × Day interactions (F_3.58,75.72_ = 2.91, p = 0.032, ηp^2^ = 0.122). The three-way Order × Diet × Day term was also significant (F_3.58,75.27_= 2.84, p = 0.035, ηp^2^ = 0.119). There was no main effect of Order (F_1,21_= 1.64, p = 0.214, ηp^2^ = 0.073). Post hoc tests were dominated by the Diet difference, with a greater effect in Order 2 mice who took a greater numbers of snacks on multiple PR days, relative to their NR counterparts. As with meals, day-to-day contrasts within Order 1 were weaker and less consistent. See Fig. S7 for sex-specific plots of number of snacks.

### 3.7. Distribution of feasts (> 5 pellets)

Our initial analysis on feasts, identified Sex as a significant factor, therefore for these analyses, Sex was included. Three-way ANOVA on the number of feasts consumed per each phase (Fig. 7A and Fig. 8A), showed a main effect of Sex (F_1,38_ = 4.45, p = 0.042), no main effect of Order (F_1,38_ = 0.014, p = 0.90) and a main effect of Diet (F_1,38_ = 18.07, p < 0.001). In addition, an interaction between Sex × Order (F_1,38_ = 4.98, p = 0.032) and Order × Diet (F_1,38_ = 6.31, p = 0.016) was detected. Post hoc comparisons indicated only male mice in Order 2, significantly increased their feast intake when switched to NR diet (p < 0.01), even though Order 2 male mice show more feasting behaviour when switched to NR diet compared with Order 1 male mice on PR diet (p < 0.05), their feast intake is not significantly higher than Order 1 NR phase (p = 0.09).

**Fig. 7.**
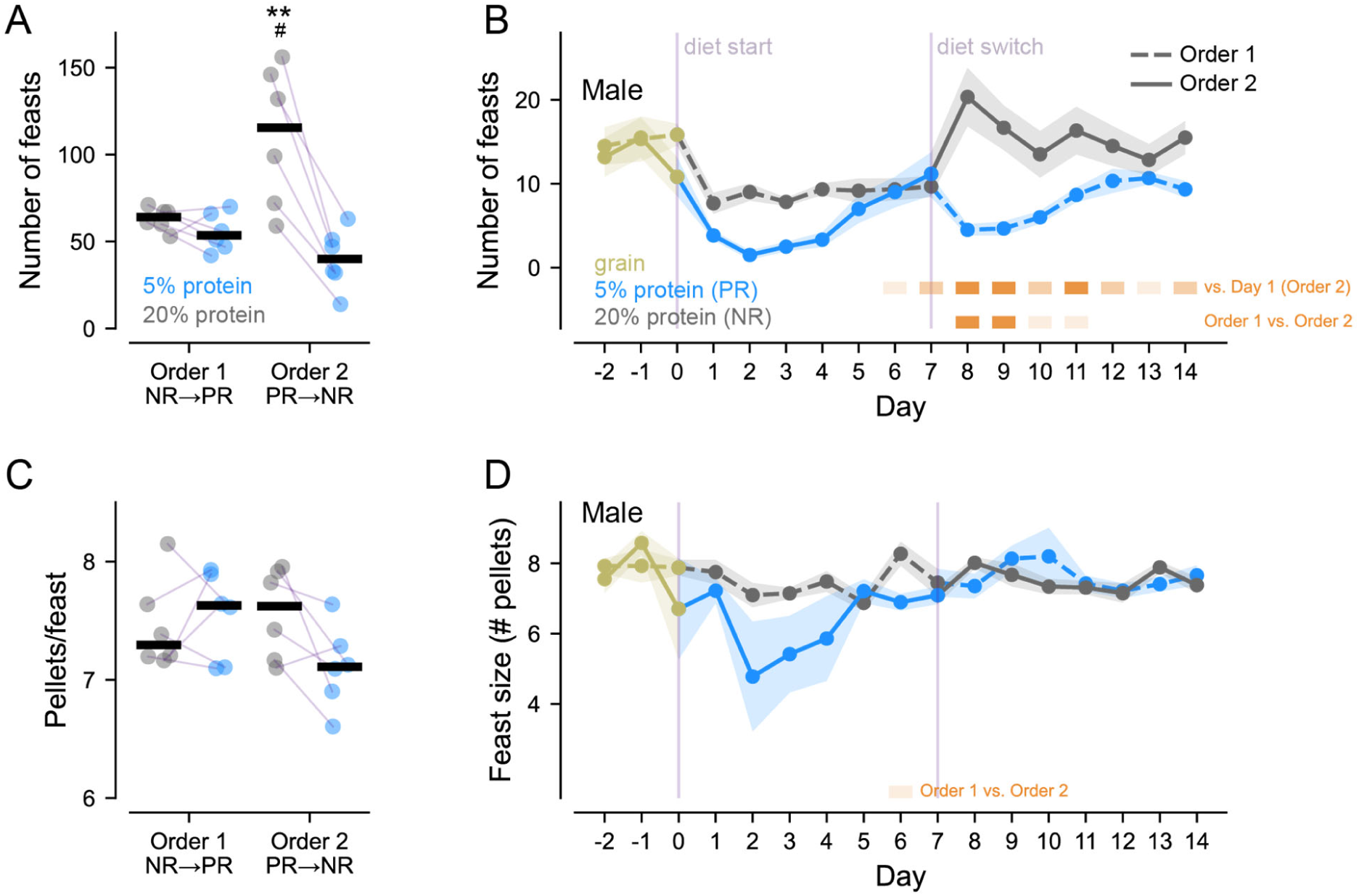
Protein manipulation alters feasting behaviour in male mice in an order-dependent manner. **(A, C)** When averaged across diet phases, male mice showed clear effects of protein restriction on feasting behaviour. Order 2 males (PR → NR) exhibited a pronounced increase in the number of feasts during the NR phase compared with their PR phase, whereas Order 1 males (NR → PR) showed no such change. **, p<0.01 vs. PR phase; #, p<0.05 vs. Order 1 mice on PR diet. Feast size remained relatively stable across diets and orders, showing only minor variability between groups. **(B, D)** Day-to-day analyses further demonstrated that Order 2 males increased feast number and frequency across the NR week, following a progressive rise during PR, whereas Order 1 males showed little daily fluctuation. Although for feast size, no consistent within-phase day-to-day pattern emerged. Statistical annotation and plotting conventions follow those described in the caption of Fig. 2.

**Fig. 8.**
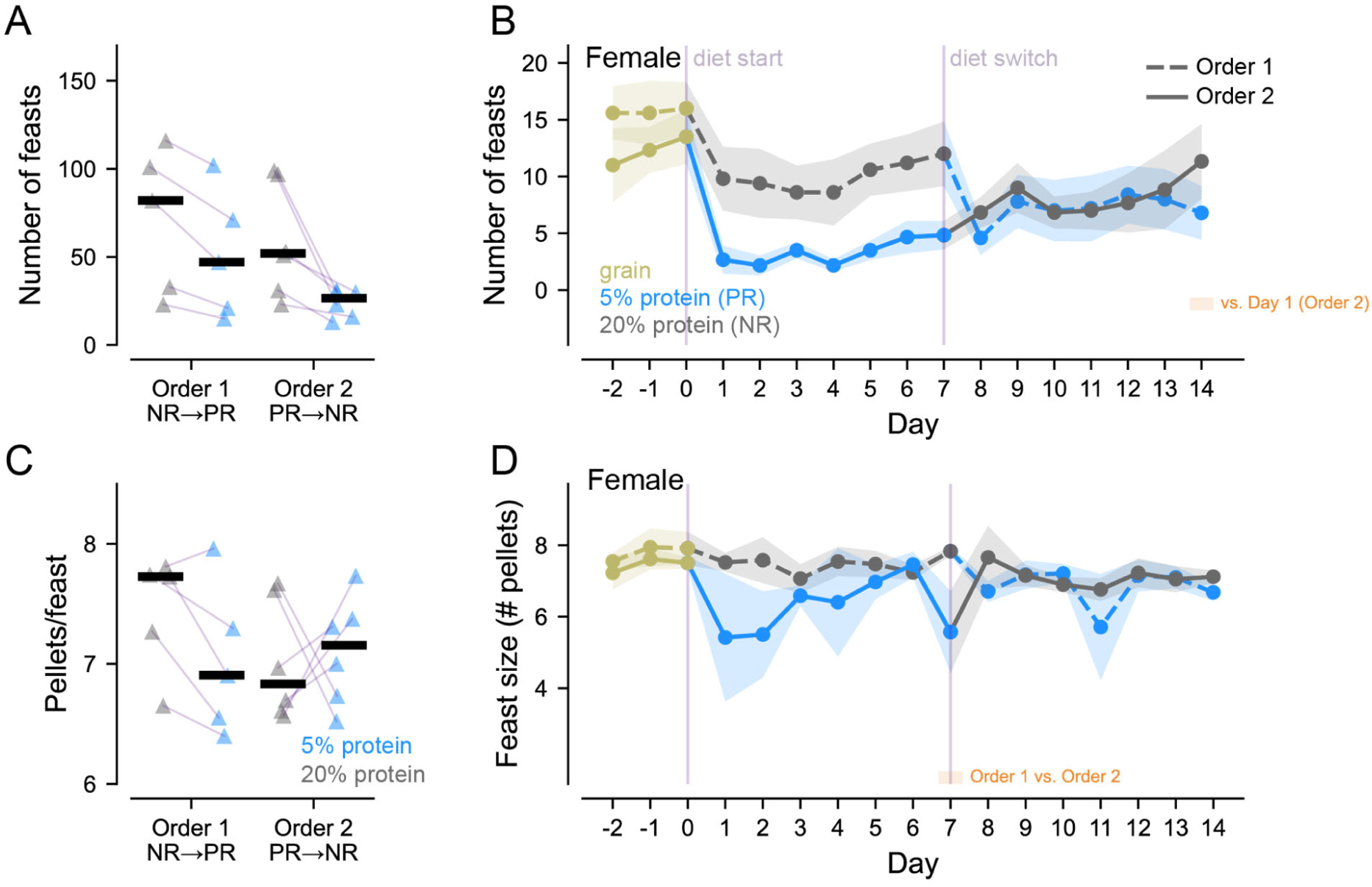
Female mice display limited alteration of feasting behaviour by dietary protein restriction. **(A)** Across diet phases, female mice showed overall fewer feasts than males, with no significant changes between NR and PR diets. While the ANOVA revealed main effects of Sex and Diet, post hoc analyses indicated that the increase in feasting under PR or NR was driven primarily by males. In females, feast size remained relatively stable across orders and diets. **(B)** Day-to-day analyses confirmed the absence of strong diet-related fluctuations in females. Feast number did not vary systematically across PR and NR weeks, and feast size remained largely constant. Thus, unlike males, female mice exhibited minimal order- or diet-dependent adaptation in feasting behaviour. Statistical annotation conventions follow those described in the caption of Fig. 2.

Analysis of feast size (Fig. 7C and Fig. 8C) revealed a main effect of Sex (F_1,38_=4.21, p=0.047), no main effect of Order (F_1,38_=1.57, p=0.217), Diet (F_1,38_=1.53, p=0.224) and no interactions other than a trend towards Sex × Order × Diet (F_1,38_=3.99, p= 0.052). No post hoc comparisons reached significance indicating that feast size is different in male and female mice but is relatively constant with respect to Diet and Order.

We examined day-to-day changes in the number of feasts across the two Diet weeks (Fig. 7B and Fig. 8B). A mixed-design repeated-measures ANOVA confirmed a large main effect of Diet (F_1,19_ = 53.33, p < 0.001, ηp^2^ = 0.737), and a strong Order × Diet interaction, (F_1,19_ = 17.77, p < 0.001, ηp^2^ = 0.483). There was also a Day effect, (F_3.12,59.22_ = 7.29, p < 0.001, ηp^2^ = 0.277), and a Diet × Day interaction, (F_2.46,46.81_ = 4.74, p = 0.009, ηp^2^ = 0.200). Critically, Diet effects depended on Sex (Order × Sex × Diet), (F_1,19_ = 7.44, p = 0.013, ηp^2^ = 0.281), and the day-wise Diet pattern differed by Sex (Sex × Diet × Day), (F_2.46,46.81_ = 5.03, p = 0.007, ηp^2^ = 0.209). The Order × Day term showed a trend (F_3.12,59.22_ = 2.27, p = 0.088, ηp^2^ = 0.107), whereas Order, Sex, and the remaining interactions were not significant (all p ≥ 0.1). Post hoc tests identified robust within-Order-2 male effects, where multiple NR days exceeded early PR, and late PR exceeded early PR. No day-to-day contrasts survived correction for Order 1 males or females in either diet phase.

On the other hand, the dynamics of feast size (Fig. 7D and Fig. 8D) showed a main effect of Order, (F_1,19_ = 6.03, p = 0.024, ηp^2^ = 0.241), and a main effect of Diet, (F_1,19_ = 12.25, p = 0.002, ηp^2^ = 0.392), accompanied by an Order × Diet interaction (F_1,19_ = 4.91, p = 0.039, ηp^2^ = 0.205). In this case, Sex and all Sex-involving terms were not significant (all ps ≥ 0.1). There was no main effect of Day, (F_3.83,72.69_ = 1.04, p = 0.391, ηp^2^ = 0.052), and no interactions with Day (all p ≥ 0.3), including the four-way term (p = 0.610), and no post hoc contrasts survived other than a single day towards the end of PR phase where Order 2 male and female mice had smaller feast size compared to Order 1 mice (ps < 0.05, day 5 and 6). Thus, feast size shows order-dependent differences (smaller in PR→NR mice) but no reliable day-to-day dynamics within diet phases.

### 3.8 Total feeding events and dynamics of interaction time with food source

To assess how overall feeding activity varies with diet phase and order of diet presentation, we conducted a two-way ANOVA on the combined count of feeding events: snacks, meals and feasts, (Fig. 9). The two-way ANOVA yielded a main effect of Order (F_1,42_ = 4.91, p = 0.032), a main effect of Diet (F_1,42_ = 16.13, p < 0.001), and a marginal interaction between Order × Diet (F_1,42_ = 3.85, p = 0.056). As such, protein-restriction was associated with more feeding events for both orders of diet presentation, and Order 2 mice exhibited more feeding events than Order 1 mice.

**Fig. 9.**
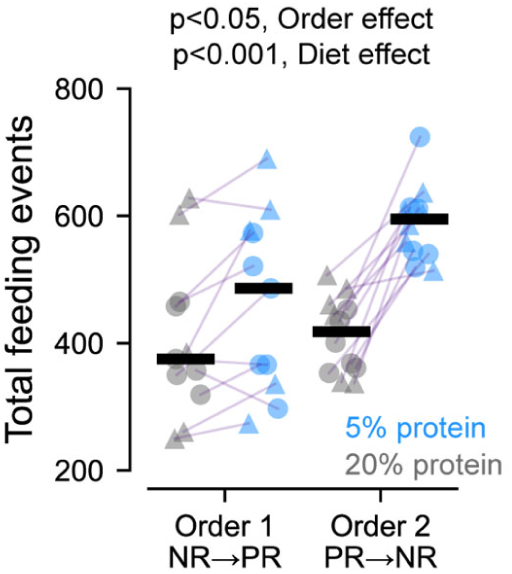
Dietary protein restriction increases total number of feeding events. When considering all feeding events combined (snacks + meals + feasts), mice on protein-restricted (PR) diet exhibited more feeding events than when on non-restricted (NR) diet. Overall, Order 2 mice exhibited more feeding events than Order 1 mice.

Subsequently, we calculated time spent interacting with FED units as a measure of exploration for food (Fig. 10), using a similar threshold of 60 s, such that clusters of logged events (pellet retrievals and nose-pokes) occurring within 60 s of one another were treated as part of the same feeding interaction bout. Importantly, nose-poke events were recorded while animals were in free-feeding mode, in which pokes had no programmed consequences and did not trigger pellet delivery; rather, they were used here as an index of interaction with the FED device and, by extension, exploration of the food source. Repeated-measures ANOVA showed a main effect of Order (F_1,21_ = 5.23, p = 0.033, ηp^2^ = 0.199), no main effect of Diet (F_1,21_ = 0.01, p = 0.932, ηp^2^ < 0.001), and a robust Day effect, (F_4.87,102.22_ = 4.74, p < 0.001, ηp^2^ = 0.184). There was a strong Diet × Day interaction, (F_3.76,78.91_ = 12.11, p < 0.001, ηp^2^ = 0.366), while Order × Day showed a trend (F_4.87,102.22_ = 2.26, p = 0.056, ηp^2^ = 0.097). The Order × Diet term was not significant, (F_1,21_ = 2.52, p = 0.127, ηp^2^ = 0.107), and the three-way Order × Diet × Day interaction was also not significant, (F_3.76,78.91_ = 1.20, p = 0.318, ηp^2^ = 0.054). Holm-corrected post hoc tests indicated that Order 2 mice, specifically at the first day on PR diet, had a significantly longer interaction time with the food source (p < 0.001) and a trend on the second day (p = 0.082) while as shown on Fig. 2B this interaction is not followed by increased food intake. This implies that the interaction is driven by increased number of nose pokes while the pellet was already in the well, reflecting greater exploration of the food source.

**Fig. 10.**
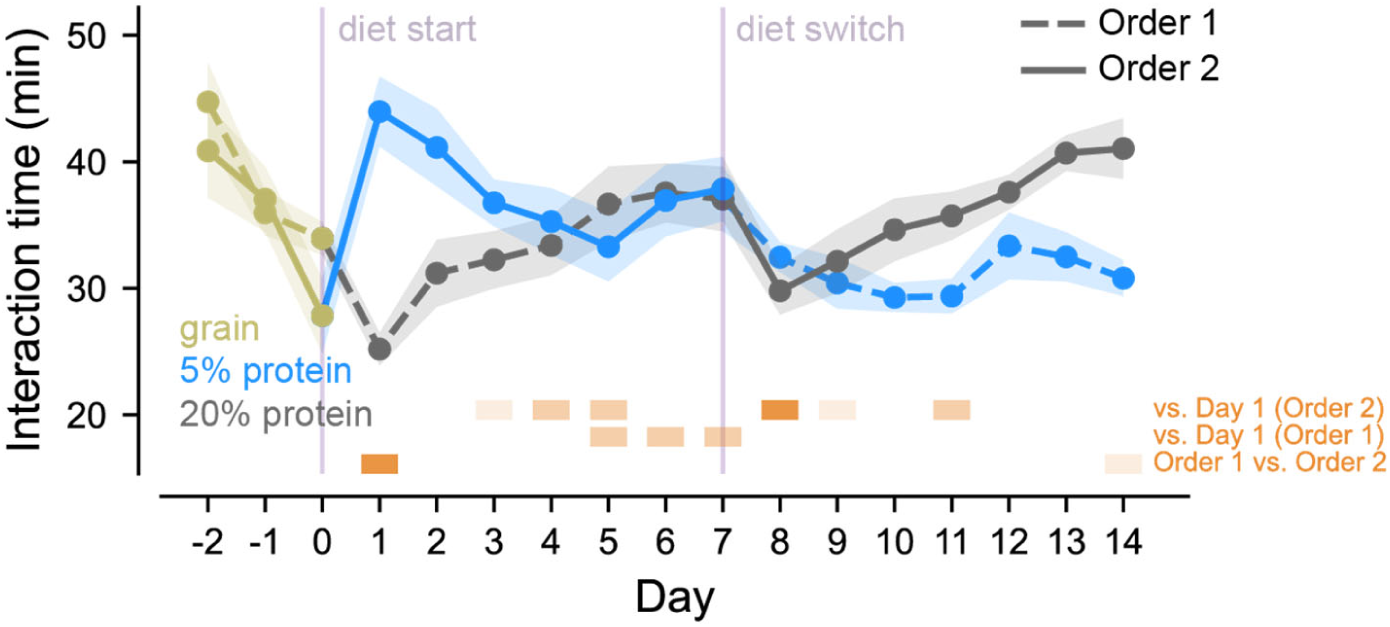
Dietary protein restriction increases exploratory interaction with the food source in an order-dependent manner. Time spent interacting with the feeding device, taken as a measure of exploratory engagement, also varied with diet phase and order. Order 2 mice spent more time interacting with FED units at the beginning of the PR phase, despite not consuming more pellets at that point, suggesting increased food-seeking or exploratory behaviour rather than consumption. Interaction time subsequently declined over days as mice adapted to the diet. Statistical annotation conventions follow those described in the caption of Fig. 2.

## 4. Discussion

Here, we show dynamic changes in feeding that occur during protein restriction in mice. We identify changes in meal patterning induced by restriction and show that effects in many cases depend on order in which mice experience each diet. Using FED3 devices to monitor feeding in the home-cage, we combined a data-driven temporal threshold, with analysis of the completed cluster-size distribution, neighbouring-size ratios and cluster-continuation probabilities, to classify eating events into snacks (1 pellet), meals (2-5 pellets), and feasts (>5 pellets). This criterion reduced some arbitrariness historically associated with many meal definitions and captured meaningful adaptations in how animals respond to changing protein availability across time.

### 4.1 Hyperphagia induced by protein-restriction

Previously, diets low in protein have been observed to induce hyperphagia in studies using animal models as well as human subjects (Gosby et al., 2014; Morrison & Laeger, 2015; Torres et al., 2022). This compensatory response in which animals are driven to eat more to defend a protein intake target is fundamental to the concept of protein leverage (Raubenheimer & Simpson, 2023; Sørensen et al., 2008). In this experiment, we did not see the expected hyperphagia induced by protein restriction when data were averaged across all days. However, examination of day-to-day intake did show evidence of restriction-induced hyperphagia emerging over time after an initial decrease in intake upon first exposure to the restricted diet. Moreover, hyperphagia persisted once mice were returned to a non-restricted diet. These dynamic changes in feeding behaviour in response to changing protein availability may be important to consider when modelling responses to protein dilute diets. In fact, a recent paper from our lab highlights that repeated alternation between a restricted and non-restricted diet leads to hyperphagia that persists across both restricted and non-restricted conditions (Volcko et al., 2024).

Interestingly, we also observed differences in intake dynamics depending on the order in which diets were presented. Mice that experienced the protein-restricted diet first (Order 2 mice), did not immediately exhibit a hyperphagic response but rather showed a gradual increase in intake beginning approximately 3 days after diet presentation. This high intake continued for several days after switching to the non-restricted diet. By contrast, mice that experienced the non-restricted diet first and only later transitioned to the restricted diet, showed relatively stable intake during the first phase, a modest transient reduction after the switch to restricted diet, and no obvious hyperphagia during the protein-restricted phase. Possibilities underlying the lack of hyperphagia include the short length (7 days) of each phase, composition of the pellets (5% protein in the restricted diet and relatively high monosaccharides in both experimental diets), the age of the mice, or interactions between these details. These details differ in other studies that have shown hyperphagia induced by low protein diets (Champeil-Potokar et al., 2021; Zaman et al., 2024) Our findings indicate that feeding responses were shaped not only by the protein content of the current diet, but also by recent dietary experience, highlighting the importance of considering how fluctuations in protein availability may impact food intake.

### 4.2 Increased exploration and promoted feeding activity

Diet composition and protein restriction can alter circadian rhythms of mice and affect the cellular machinery involved in regulation of feeding (Cheng et al., 2021; Han et al., 2024). Here, with free access to food, we did not identify changes in feeding activity such as a shift in active feeding to the light phase. Rather, increased feeding occurred during the expected peak hours in the dark phase. With respect to interaction time with FEDs, we observed an increase in interactions during initial exposure to the protein-restricted diet, despite this coinciding with a reduction in pellet retrievals. In addition, summing up total feeding activities per phase, confirmed that all mice exhibited more feeding events during the phase with restricted diet. These observations suggest that protein-restricted mice continued to explore the food source, potentially in search of a pellet with desired nutritional value. Such increased interaction with the feeding source may reflect foraging-like, appetitive behaviour rather than consummatory intake itself. Exploratory food-seeking behaviours are known to increase under nutrient-deficient states, and studies examining foraging more directly have shown that protein restriction can increase horizontal and vertical exploration in mice, consistent with greater motivation to seek nutritionally appropriate food (Heinz et al., 2021; Keen-Rhinehart & Bartness, 2007; S. Wu et al., 2023).

### 4.3 Defining feeding events and analysis of meal patterning

Meal definition is pivotal to the analysis of meal patterning and understanding regulation of food intake, yet a validated and unified standard remains elusive. Various criteria have been suggested to delineate the definition of meals, often focusing on the intervals between feeding events or quantity of food ingested (Ali & Kravitz, 2018). In rat studies, various temporal thresholds have been used, including a 40 min gap threshold proposed by Le Magnen & Devos (1980) and a 10-minute inter-meal interval defined through log-survivorship analysis (Castonguay et al., 1986). An alternative approach was taken by Zorrilla et al. (2005) by incorporating water intake into the meal definition and identifying a 5 min breakpoint between feeding or drinking events of rats. Kanoski and colleagues defined a meal in rats as bouts of feeding separated by at least 10 min gap, with a minimum intake of 0.25 g (Kanoski et al., 2012), while Martire and colleagues used behavioural criteria to differentiate meals and snacks based on the sequence of behaviours following a feeding event (Martire et al., 2013). Research using systems that monitor weight of food (e.g., BioDAQ) defined a meal as any feeding episode containing at least 0.2 g of food, isolated by an inter-meal interval of 5 min (Boyle et al., 2012). Studies in mice have used similar strategies, with one group defining a meal as at least 0.02 g of food consumed (Goebel et al., 2011), and another stipulating 0.05 g or more within a 15 min window (Rathod & Di Fulvio, 2021; Tabarin et al., 2007). The first study using FED3 units classified pellets eaten within 1 min of each other as a meal if they totalled at least 0.1 g (five 20 mg pellets) (Matikainen-Ankney et al., 2021), while the reasons for the choice of 0.1 g as a threshold was not specified.

Here, we employed a mathematical approach and combined the classic concept of a temporal threshold with a criterion to demarcate different types of feeding bout. Our method identified a natural temporal breakpoint at α_IPI_ = 60 s, similar to that suggested by Matikainen-Ankney et al. (2021). As such, most feeding events are composed of short trains of pellet retrievals, punctuated by brief pauses, while prolonged breaks (>60 s) signal termination of the feeding events. Following the 60 s temporal threshold, examination of the completed cluster-size distributions showed that 2-5 pellet events comprised the predominant multi-pellet feeding range, whereas events containing more than five pellets formed a less frequent extended range. Neighboring completed-cluster ratios highlighted a reduction between five- and six-pellet events, while cluster-continuation probabilities showed gradual progression across cluster sizes rather than an abrupt transition. Together, these complementary analyses supported the use of five pellets as an operational upper boundary of the predominant meal range, allowing feeding events to be classified into snacks (1 pellet), meals (2– 5 pellets), and feasts (>5 pellets). Interestingly, in the reference group we used to categorise these events (mice eating non-restricted pellets without previous exposure to protein-restriction), we found the separation between meals and feasts to be more pronounced in male than in female mice (Fig. S8). Future work should collect more data over longer periods to determine whether there are consistent sex-specific effects on feeding microstructure.

Using these definitions across all conditions, we found that protein restriction increased total meal number (and hence, meal frequency), while reducing meal size. In addition, we observed that restriction also increased the number of snacks. When mice switched back to the non-restricted diet, the balance among these components changed markedly, such that meal size increased, snacking declined, and a “feasting” behaviour emerged. This shift in feeding pattern likely reflects an adaptive response to optimise nutrient acquisition and maintain energy balance under conditions of protein scarcity.

### 4.4 Potential mechanisms underlying behavioural dynamics

Future work will be needed to understand the hormonal and neural mechanisms that underlie these changes in intake and meal patterning. One important factor could be ghrelin dynamics, as we previously showed that protein restriction elevates circulating ghrelin, and that this elevation can persist even after normal dietary protein is restored (Volcko et al., 2024). Ghrelin is best known for increasing during caloric insufficiency and acts in several brain regions to stimulate appetite. Importantly, its orexigenic effect does not immediately normalise when food availability returns (Fernandez et al., 2018), thus it could underlie the increased intake we observed once the non-restricted diet became available. Interactions with signals that relay protein insufficiency (e.g., FGF21) (Hill et al., 2019; Laeger et al., 2014) and conventional satiety signals induced by protein ingestion (e.g., GLP-1, CCK, and PYY) (Batterham et al., 2006; Morrison et al., 2012; Morrison & Laeger, 2015; van der Klaauw et al., 2013) are also likely to be involved. For FGF21, the dual effects of increasing protein intake while decreasing sweet preference (e.g., (von Holstein-Rathlou et al., 2016; C.-T. Wu et al., 2024)) might be especially important for understanding the switch between generalised hyperphagia when dietary protein is moderately low to protein specific appetite without pronounced hyperphagia when dietary protein is further reduced.

As well as the hypothalamic and hindbrain circuits engaged by these hormones, reward circuitry is also strongly implicated in protein-restriction induced behavioural changes. Protein restriction increases ex vivo dopamine release (Naneix et al., 2020) and leads to increased ventral tegmental area activity in response to protein consumption in vivo (Chiacchierini et al., 2021). Such restriction induced shifts in dopamine responses to macronutrients appear to be FGF21-dependent (Khan et al., 2025). Furthermore protein restriction diminishes the reward value of non-protein foods as prolonged restriction blunts nucleus accumbens dopamine release induced by sucrose intake and reduces behavioural preference for sweet foods (C.-T. Wu et al., 2024), implying a transient re-allocation of reward processing toward protein. Collectively, these findings support the concept of a dynamic, state-dependent “protein appetite”, whereby reward circuits promote the acquisition and overconsumption of protein when the organism has been in a protein-deficient state.

### 4.5 Limitations and future directions

An important limitation of FED devices (as well as most other food monitoring devices, such as Bio-DAQ) is that “pellet retrieval” is logged, but not “pellet intake”. Thus, one should consider the possibility of mice collecting and discarding pellets, which might have influenced the detailed analysis of meal microstructure. Nevertheless, ongoing research in our lab using video recording coupled to pellet delivery recordings shows that >95% of the pellets retrieved are also eaten.

The order effects in our experiment, suggest history-dependence, inviting designs with washout or counter-balanced multi-phase schedules (e.g., PR↔NR↔PR) and choice tests to track evolving macronutrient preference alongside meal structure. In this study, the habituation time used before diet manipulations was also relatively short and discrimination of order effects may have been enhanced with a longer habituation and a more stable baseline. Initial effects on pellet intake due to potential neophobia should not be ignored although we have collected pilot data showing that naïve mice will readily consume both types of pellets in short (e.g., 15 min) tests. In addition, it should be noted that the mice in this study were young adults (7-9 weeks at start) and it will be important to test whether the same effects are observed in older cohorts. While our interaction-time metric indicates exploration/food-seeking, pairing FED3 logs with concurrent video recordings (e.g., to disambiguate poke patterns and pellet approach) would test whether exploratory surges precede behavioural shifts. We recently developed a solution that can be paired with FEDs to easily make such paired recordings (Taghipourbibalan & McCutcheon, 2026).

Feeding events were classified in this study by examining data from the non-restricted phase of Order 1 mice as a reference. It is possible that this classification – particularly between meals and feasts - could differ based on experimental parameters such as age of mice, body weight, and pellet composition, and sex. Indeed, Fig. S8 suggests that patterns of eating events may differ between male and female mice. As such, our findings here could be confirmed and made more robust by considering longer periods of data collection under stable conditions to determine how consistent the boundary between meals and feasts is.

Targeted endocrine assays, particularly of FGF21, would give insight into the contribution of underlying hormonal changes. This could be especially interesting given FGF21’s role not just in protein appetite but reduction in sweet preference. It is of note that the experimental diets were high in monosaccharides and thus increases in FGF21 could be affecting intake and limiting hyperphagic responses. In addition, adding neural readouts (e.g., VTA activity during protein vs. carbohydrate bouts) across PR→NR transitions could provide a mechanism that underpins the observed snack→ meal→ feast reconfiguration based on concepts such as changing stimulus evaluation and effort requirements. Finally, given sex- and order-specific weight effects, powering sex-stratified cohorts and body-composition measurements would refine interpretations of intake-weight coupling.

## Author contributions

HT: Conceptualization, Formal analysis, Investigation, Visualization, Writing – original draft. JM: Conceptualization, Funding acquisition, Supervision, Visualization, Writing – review and editing.

## Acknowledgments

The authors would like to acknowledge the staff at the Department of Comparative Medicine (AKM) and the associated animal facility of the Faculty of Health Sciences at UiT and the staff working in the workshops of the Faculty of Health Sciences at UiT.

## References

Ali, M. A., & Kravitz, A. V. (2018). Challenges in quantifying food intake in rodents. Brain Research, 1693, 188–191. 10.1016/j.brainres.2018.02.040

Barrett, M. R., Pan, Y., Murrell, C. L., Karolczak, E. O., Wang, J., Fang, L. Z., Thompson, J. M., Chang, Y.-H., Casey, E., Czarny, J. E., So, W. L., Reichenbach, A., Stark, R., Taghipourbibalan, H., Penna, S. R., McCullough, K. B., Westbrook, S. R., Chatterjee Basu, G., Matikainen-Ankney, B., … Kravitz, A. V. (2025). A simple action reduces high-fat diet intake and obesity in mice. Current Biology, S0960982225006839. 10.1016/j.cub.2025.05.067

Batterham, R. L., Heffron, H., Kapoor, S., Chivers, J. E., Chandarana, K., Herzog, H., Le Roux, C. W., Thomas, E. L., Bell, J. D., & Withers, D. J. (2006). Critical role for peptide YY in protein-mediated satiation and body-weight regulation. Cell Metabolism, 4(3), 223–233. 10.1016/j.cmet.2006.08.001

Boyle, C. N., Lorenzen, S. M., Compton, D., & Watts, A. G. (2012). Dehydration-Anorexia Derives From A Reduction In Meal Size, But Not Meal Number. Physiology & Behavior, 105(2), 305–314. 10.1016/j.physbeh.2011.08.005

Castonguay, T. W., Kaiser, L. L., & Stern, J. S. (1986). Meal pattern analysis: Artifacts, assumptions and implications. Brain Research Bulletin, 17(3), 439–443. 10.1016/0361-9230(86)90252-2

Champeil-Potokar, G., Crossouard, L., Jérôme, N., Ouali, C., Darcel, N., Davidenko, O., Rampin, O., Bombail, V., & Denis, I. (2021). Diet Protein Content and Individual Phenotype Affect Food Intake and Protein Appetence in Rats. The Journal of Nutrition, 151(5), 1311–1319. 10.1093/jn/nxaa455

Cheng, H., Liu, Z., Wu, G., Ho, C.-T., Li, D., & Xie, Z. (2021). Dietary compounds regulating the mammal peripheral circadian rhythms and modulating metabolic outcomes. Journal of Functional Foods, 78, 104370. 10.1016/j.jff.2021.104370

Chiacchierini, G., Naneix, F., Peters, K. Z., Apergis-Schoute, J., Snoeren, E. M. S., & McCutcheon, J. E. (2021). Protein appetite drives macronutrient-related differences in ventral tegmental area neural activity. Journal of Neuroscience, 41(23), 5080–5092. 10.1523/JNEUROSCI.3082-20.2021

Cohen, L. R., & Woodside, B. C. (1989). Self-selection of protein during pregnancy and lactation in rats. Appetite, 12(2), 119–136. 10.1016/0195-6663(89)90101-3

Fernandez, G., Cabral, A., Andreoli, M. F., Labarthe, A., M’Kadmi, C., Ramos, J. G., Marie, J., Fehrentz, J.-A., Epelbaum, J., Tolle, V., & Perello, M. (2018). Evidence Supporting a Role for Constitutive Ghrelin Receptor Signaling in Fasting-Induced Hyperphagia in Male Mice. Endocrinology, 159(2), 1021–1034. 10.1210/en.2017-03101

Goebel, M., Stengel, A., Wang, L., & Taché, Y. (2011). Central nesfatin-1 reduces the nocturnal food intake in mice by reducing meal size and increasing inter-meal intervals. Peptides, 32(1), 36–43. 10.1016/j.peptides.2010.09.027

Gosby, A. K., Conigrave, A. D., Raubenheimer, D., & Simpson, S. J. (2014). Protein leverage and energy intake. Obesity Reviews, 15(3), 183–191. 10.1111/obr.12131

Han, Y., Shon, J., Kwon, S. Y., & Park, Y. J. (2024). Effects of Dietary Protein Intake Levels on Peripheral Circadian Rhythm in Mice. International Journal of Molecular Sciences, 25(13), 7373. 10.3390/ijms25137373

Heinz, D. E., Schöttle, V. A., Nemcova, P., Binder, F. P., Ebert, T., Domschke, K., & Wotjak, C. T. (2021). Exploratory drive, fear, and anxiety are dissociable and independent components in foraging mice. Translational Psychiatry, 11(1), 1–12. 10.1038/s41398-021-01458-9

Hill, C. M., Laeger, T., Dehner, M., Albarado, D. C., Clarke, B., Wanders, D., Burke, S. J., Collier, J. J., Qualls-Creekmore, E., Solon-Biet, S. M., Simpson, S. J., Berthoud, H.-R., Münzberg, H., & Morrison, C. D. (2019). FGF21 Signals Protein Status to the Brain and Adaptively Regulates Food Choice and Metabolism. Cell Reports, 27(10), 2934–2947.e3. 10.1016/j.celrep.2019.05.022

Kanoski, S. E., Zhao, S., Guarnieri, D. J., DiLeone, R. J., Yan, J., De Jonghe, B. C., Bence, K. K., Hayes, M. R., & Grill, H. J. (2012). Endogenous leptin receptor signaling in the medial nucleus tractus solitarius affects meal size and potentiates intestinal satiation signals. American Journal of Physiology - Endocrinology and Metabolism, 303(4), E496–E503. 10.1152/ajpendo.00205.2012

Keen-Rhinehart, E., & Bartness, T. J. (2007). NPY Y1 receptor is involved in ghrelin- and fasting-induced increases in foraging, food hoarding, and food intake. *American Journal of Physiology - Regulatory*, Integrative and Comparative Physiology, 292(4), R1728–R1737. 10.1152/ajpregu.00597.2006

Khan, M. S. H., Kim, S. Q., Ross, R. C., Corpodean, F., Spann, R. A., Albarado, D. A., Fernandez-Kim, S. O., Clarke, B., Berthoud, H.-R., Münzberg, H., McDougal, D. H., He, Y., Yu, S., Albaugh, V. L., Soto, P. L., & Morrison, C. D. (2025). FGF21 acts in the brain to drive macronutrient-specific changes in behavioral motivation and brain reward signaling. Molecular Metabolism, 91, 102068. 10.1016/j.molmet.2024.102068

Laeger, T., Henagan, T. M., Albarado, D. C., Redman, L. M., Bray, G. A., Noland, R. C., Münzberg, H., Hutson, S. M., Gettys, T. W., Schwartz, M. W., & Morrison, C. D. (2014). FGF21 is an endocrine signal of protein restriction. The Journal of Clinical Investigation, 124(9), 3913. 10.1172/JCI74915

Le Magnen, J., & Devos, M. (1980). Parameters of the meal pattern in rats: Their assessment and physiological significance. Neuroscience & Biobehavioral Reviews, 4, 1–11. 10.1016/0149-7634(80)90040-8

Martire, S. I., Holmes, N., Westbrook, R. F., & Morris, M. J. (2013). Altered Feeding Patterns in Rats Exposed to a Palatable Cafeteria Diet: Increased Snacking and Its Implications for Development of Obesity. PLoS ONE, 8(4), e60407. 10.1371/journal.pone.0060407

Matikainen-Ankney, B. A., Earnest, T., Ali, M., Casey, E., Wang, J. G., Sutton, A. K., Legaria, A. A., Barclay, K. M., Murdaugh, L. B., Norris, M. R., Chang, Y.-H., Nguyen, K. P., Lin, E., Reichenbach, A., Clarke, R. E., Stark, R., Conway, S. M., Carvalho, F., Al-Hasani, R., … Kravitz, A. V. (2021). An open-source device for measuring food intake and operant behavior in rodent home-cages. eLife, 10, e66173. 10.7554/eLife.66173

Morrison, C. D., & Laeger, T. (2015). Protein-dependent regulation of feeding and metabolism. Trends in Endocrinology and Metabolism: TEM, 26(5), 256–262. 10.1016/j.tem.2015.02.008

Morrison, C. D., Reed, S. D., & Henagan, T. M. (2012). Homeostatic regulation of protein intake: In search of a mechanism. American Journal of Physiology - Regulatory Integrative and Comparative Physiology, 302(8), 917–928. 10.1152/ajpregu.00609.2011

Murphy, M., Peters, K. Z., Denton, B. S., Lee, K. A., Chadchankar, H., & McCutcheon, J. E. (2018). Restriction of dietary protein leads to conditioned protein preference and elevated palatability of protein-containing food in rats. Physiology & Behavior, 184, 235–241. 10.1016/J.PHYSBEH.2017.12.011

Musten, B., Peace, D., & Anderson, G. H. (1974). Food Intake Regulation in the Weanling Rat: Self-selection of Protein and Energy. The Journal of Nutrition, 104(5), 563–572. 10.1093/JN/104.5.563

Naneix, F., Peters, K. Z., Young, A. M. J., & McCutcheon, J. E. (2020). Age-dependent effects of protein restriction on dopamine release. Neuropsychopharmacology 2020 46:2, 46(2), 394–403. 10.1038/s41386-020-0783-z

Qasem, R. J., Li, J., Tang, H. M., Pontiggia, L., & D’mello, A. P. (2016). Maternal protein restriction during pregnancy and lactation alters central leptin signalling, increases food intake, and decreases bone mass in 1 year old rat offspring. Clinical and Experimental Pharmacology and Physiology, 43(4), 494–502. 10.1111/1440-1681.12545

Rathod, Y. D., & Di Fulvio, M. (2021). The feeding microstructure of male and female mice. PLOS ONE, 16(2), e0246569. 10.1371/JOURNAL.PONE.0246569

Raubenheimer, D., & Simpson, S. J. (2023). Protein appetite as an integrator in the obesity system: The protein leverage hypothesis. Philosophical Transactions of the Royal Society B, 378(1888), 20220212. 10.1098/RSTB.2022.0212

Simpson, S. J., & Raubenheimer, D. (2005). Obesity: The protein leverage hypothesis. Obesity Reviews, 6(2), 133–142. 10.1111/j.1467-789X.2005.00178.x

Sørensen, A., Mayntz, D., Raubenheimer, D., & Simpson, S. (2008). Protein-leverage in mice: The geometry of macronutrient balancing and consequences for fat deposition. *Obesity (Silver Spring*, Md.), 16(3), 566–571. 10.1038/OBY.2007.58

Tabarin, A., Diz-Chaves, Y., Consoli, D., Monsaingeon, M., Bale, T. L., Culler, M. D., Datta, R., Drago, F., Vale, W. W., Koob, G. F., Zorrilla, E. P., & Contarino, A. (2007). Role of the corticotropin-releasing factor receptor type 2 in the control of food intake in mice: A meal pattern analysis. European Journal of Neuroscience, 26(8), 2303–2314. 10.1111/j.1460-9568.2007.05856.x

Taghipourbibalan, H., & McCutcheon, J. E. (2026). RTFED, an open-source versatile tool for home-cage monitoring of behaviour and fibre photometry recording in mice. Journal of Neuroscience Methods, 425, 110604. 10.1016/j.jneumeth.2025.110604

Torres, F., Khan, S., Fernandez-Kim, S. O., Spann, R., Albarado, D., Wagner, T. J., Morrison, C. D., & Soto, P. L. (2022). Dynamic effects of dietary protein restriction on body weights, food consumption, and protein preference in C57BL6/J and Fgf21-KO mice. Journal of the Experimental Analysis of Behavior, 117(3), 346–362. 10.1002/jeab.745

van der Klaauw, A. A., Keogh, J. M., Henning, E., Trowse, V. M., Dhillo, W. S., Ghatei, M. A., & Farooqi, I. S. (2013). High Protein Intake Stimulates Postprandial GLP1 and PYY Release. Obesity (Silver Spring, Md.), 21(8), 1602–1607. 10.1002/oby.20154

Volcko, K. L., & McCutcheon, J. E. (2022). Protein preference and elevated plasma FGF21 induced by dietary protein restriction is similar in both male and female mice. Physiology & Behavior, 257, 113994. 10.1016/J.PHYSBEH.2022.113994

Volcko, K. L., Taghipourbibalan, H., & McCutcheon, J. E. (2024). Intermittent protein restriction elevates food intake and plasma ghrelin in male mice. Appetite, 203, 107671. 10.1016/j.appet.2024.107671

von Holstein-Rathlou, S., BonDurant, L., Peltekian, L., Naber, M. C., Yin, T. C., Claflin, K. E., Urizar, A. I., Madsen, A. N., Ratner, C., Holst, B., Karstoft, K., Vandenbeuch, A., Anderson, C. B., Cassell, M. D., Thompson, A. P., Solomon, T. P. J., Rahmouni, K., Kinnamon, S. C., Pieper, A. A., … Potthoff, M. J. (2016). FGF21 Mediates Endocrine Control of Simple Sugar Intake and Sweet Taste Preference by the Liver. Cell Metabolism, 23(2), 335–343. 10.1016/j.cmet.2015.12.003

Watts, A. G., Kanoski, S. E., Sanchez-Watts, G., & Langhans, W. (2022). THE PHYSIOLOGICAL CONTROL OF EATING: SIGNALS, NEURONS, AND NETWORKS. Physiological Reviews, 102(2), 689–813. 10.1152/PHYSREV.00028.2020/ASSET/IMAGES/LARGE/PHYSREV.00028.2020_F022.JPEG

Wu, C.-T., Gonzalez Magaña, D., Roshgadol, J., Tian, L., & Ryan, K. K. (2024). Dietary protein restriction diminishes sucrose reward and reduces sucrose-evoked mesolimbic dopamine signaling in mice. Appetite, 203, 107673. 10.1016/j.appet.2024.107673

Wu, S., Wang, J., Xu, Y., Zhang, Z., Jin, X., Liang, Y., Ge, Y., Zhan, H., Peng, L., Luo, D., Li, M., Bi, W., Guan, Q., & He, Z. (2023). Energy deficiency promotes rhythmic foraging behavior by activating neurons in paraventricular hypothalamic nucleus. Frontiers in Nutrition, 10. 10.3389/fnut.2023.1278906

Yap, Y. W., Rusu, P. M., Chan, A. Y., Fam, B. C., Jungmann, A., Solon-Biet, S. M., Barlow, C. K., Creek, D. J., Huang, C., Schittenhelm, R. B., Morgan, B., Schmoll, D., Kiens, B., Piper, M. D. W., Heikenwälder, M., Simpson, S. J., Bröer, S., Andrikopoulos, S., Müller, O. J., & Rose, A. J. (2020). Restriction of essential amino acids dictates the systemic metabolic response to dietary protein dilution. Nature Communications, 11(1), 2894. 10.1038/s41467-020-16568-z

Zaman, K., Mun, H., Solon-Biet, S. M., Senior, A. M., Raubenheimer, D., Simpson, S. J., & Conigrave, A. D. (2024). Mice Regulate Dietary Amino Acid Balance and Energy Intake by Selecting between Complementary Protein Sources. The Journal of Nutrition, 154(6), 1766–1780. 10.1016/j.tjnut.2024.04.007

Zheng, J., Zhang, L., Liu, J., Li, Y., & Zhang, J. (2021). Long-Term Effects of Maternal Low-Protein Diet and Post-weaning High-Fat Feeding on Glucose Metabolism and Hypothalamic POMC Promoter Methylation in Offspring Mice. Frontiers in Nutrition, 8(August), 1–9. 10.3389/fnut.2021.657848

Zorrilla, E. P., Inoue, K., Fekete, É. M., Tabarin, A., Valdez, G. R., & Koob, G. F. (2005). Measuring meals: Structure of prandial food and water intake of rats. *American Journal of Physiology-Regulatory*, Integrative and Comparative Physiology, 288(6), R1450–R1467. 10.1152/ajpregu.00175.2004

