## Supplementary Material for "Dynamics of feeding behaviour and meal patterning in protein-restricted mice"

Contents

**Table S1. Experimental diets used in this study** **2**

**2.4.S1. A comprehensive list of R packages, statistical functions, and analytical workflows used** **3**

**Fig. S1. Dynamics of food intake separated by sex** **5**

**Fig. S2. Caloric intake during diet transition** **5**

**3.4.S1. Detailed definition of feeding microstructures using interpellet intervals and cluster-continuation analysis** **7**

Fig. S3. KDE plot of interpellet intervals . . . . . . . . . . . . . . . . . . . . . . . . 7

Fig. S4. Feeding-event cluster sizes and cluster-continuation probability . . . . . . . 11

**Fig. S5. Dynamics of numbers of meals separated by sex** **12**

**Fig. S6. Dynamics of size of meals separated by sex** **13**

**Fig. S7. Dynamics of snack numbers separated by sex** **14**

**Fig. S8. Clusters separated by sex** **15**

Table. S1. Experimental diets used in this study. Ingredients and macronutrient breakdown in control (20% casein) and protein-restricted diet (5% casein).

| Ingredient | Gram (20% Casein) | Gram/kg (20% Casein) | Gram (5% Casein) | Gram/kg (5% Casein) |
|---|---|---|---|---|
| Casein | 200 | 199.76 | 50 | 50.774 |
| L-Cystine | 3 | 2.996 | 0.75 | 0.762 |
| Dextrose | 500.7 | 482.6 | 635 | 627.331 |
| Sucrose | 107.0777 | 106.949 | 107.077 | 108.735 |
| Cellulose | 50 | 49.94 | 50 | 50.774 |
| Soybean Oil | 25 | 24.97 | 25 | 25.387 |
| Hydrogenated Cottonseed Oil | 75 | 74.91 | 75 | 76.161 |
| Mineral Mix (S10022C) | 3.5 | 3.496 | 3.5 | 3.554 |
| Calcium Carbonate | 12.495 | 12.48 | 8.7 | 8.835 |
| Calcium Phosphate Dibasic | 0 | 0 | 5.3 | 5.382 |
| Potassium Citrate | 2.4773 | 2.474 | 2.4773 | 2.516 |
| Potassium Phosphate Monobasic | 6.86 | 6.852 | 6.86 | 6.966 |
| Sodium Chloride | 2.59 | 2.587 | 2.59 | 2.63 |
| Vitamin Mix (AIN-93) | 10 | 9.988 | 10 | 10.155 |
| Choline Bitartrate | 2.5 | 2.497 | 2.5 | 2.539 |
| Tableting Aids | 17.5 | 17.5 | 17.5 | 17.5 |
| **Total** | 1001.2 | 1000 | 984.75 | 1000 |

| Nutrient | gram% (20% Casein) | kcal/gram (20% Casein) | %kcal (20% Casein) | gram% (5% Casein) | kcal/gram (5% Casein) | %kcal (5% Casein) |
|---|---|---|---|---|---|---|
| Protein | 18 | 0.72 | 18 | 5 | 0.18 | 5 |
| Fat | 10 | 0.9 | 22 | 10 | 0.9 | 23 |
| Carbohydrate | 60 | 2.39 | 60 | 73 | 2.91 | 73 |
| | | 4.01 | 100 | | 3.99 | 100 |

### 2.4.S1. A comprehensive list of R packages, statistical functions, and analytical workflows used

### R packages used for statistical analysis

The following R packages were used for data wrangling and statistical analysis:

- **tidyverse** for data manipulation and reshaping.
- **afex** and **car** for repeated-measures ANOVA and Type II/III ANOVA.
- **emmeans** and **multcomp** for post hoc pairwise comparisons, with Holm corrections as the final reported method.
- **lme4** and **lmerTest** for fitting mixed-effects models to account for repeated measures by individual mice.

#### 1. Time-course line-plot data

For measures collected across multiple days, including pellet intake, meal frequency and snack frequency over time:

- Data were reshaped to a long format, with each time point treated as a repeated measure within each mouse.
- Repeated-measures ANOVA was conducted using `aov_car()` from `afex` to test main and interaction effects of diet phase, sex, treatment order and time.
- Where significant effects were found, pairwise post hoc comparisons were conducted with Holm correction for multiple comparisons, for example via `emmeans` or `pairwise.t.test`.
- Tukey's HSD was also computed for exploratory purposes, but only Holm-adjusted *p*-values are reported in the manuscript.

#### 2. Hourly data, heatmaps and mixed-effects models

For hourly feeding data used to create heatmaps:

- Data comprised repeated observations for each mouse across different hours of the day.
- A linear mixed-effects model, using `lmer()`, was fit with fixed effects for diet, sex, order and hour, and a random intercept for each mouse to account for within-subject correlation.
- Type III ANOVA on the fitted model, via `anova()` from `lmerTest`, was used to assess the significance of main effects and interactions.
- Holm-adjusted post hoc tests via `emmeans` were used to further explore significant interactions at specific hour blocks.

#### 3. Scatter-plot metrics

For data summarised by phase or overall, including total pellets per phase, body weight per phase, feeding-event frequency and feeding-event size:

- Data were converted to a long format with PR vs. NR as within-subject factors where appropriate.
- Depending on the number of factors, one-, two- or three-way ANOVA was performed using `Anova()` from the `car` package with Type II sums of squares.
- For repeated measures or paired designs, an appropriate within-subject or blocking factor was included, or a repeated-measures design was specified in `aov_car()`.
- Holm-adjusted post hoc tests were performed to detect pairwise group differences. Tukey's HSD was also computed, but only Holm-corrected results are reported.

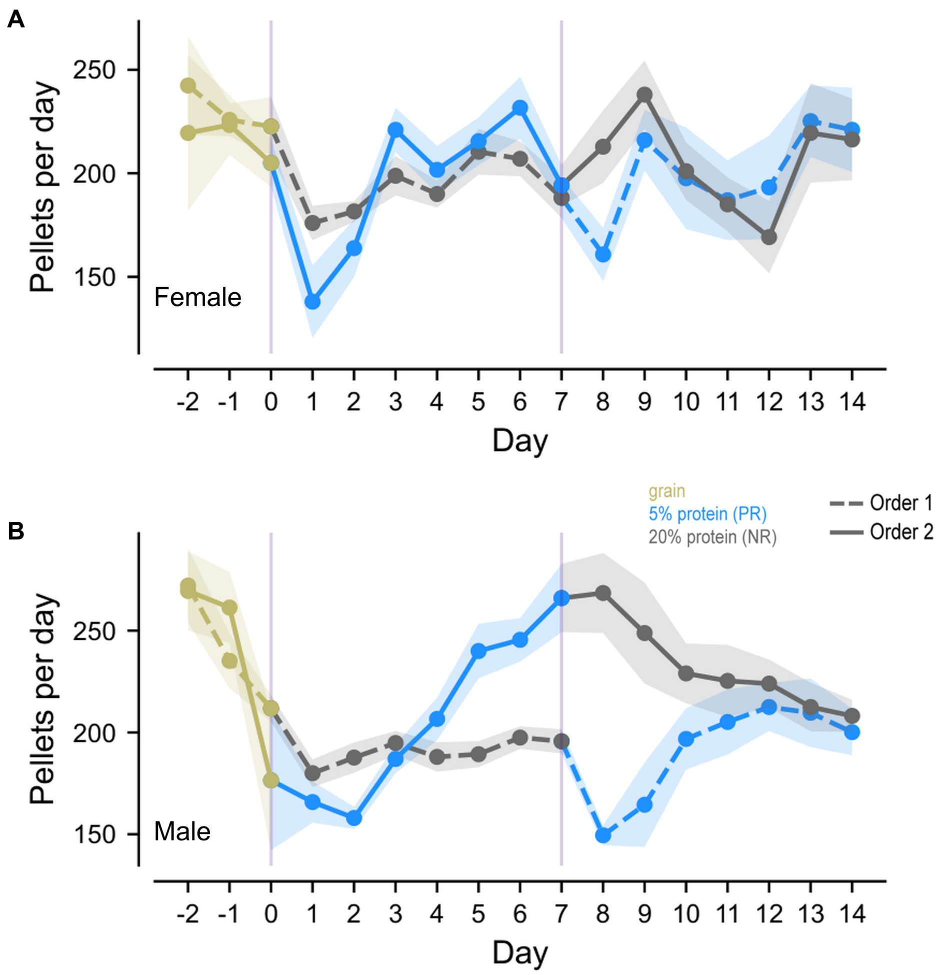


Fig. S1. Day-to-day pellet intake shown separately for female (A) and male (B) mice. Data are presented as mean ± SEM. Dashed lines indicate Order 1 and solid lines indicate Order 2. Colours denote diet (grain, 5% protein (PR), and 20% protein (NR)). These sex-specific plots correspond to the pooled analysis shown in Fig. 2B.

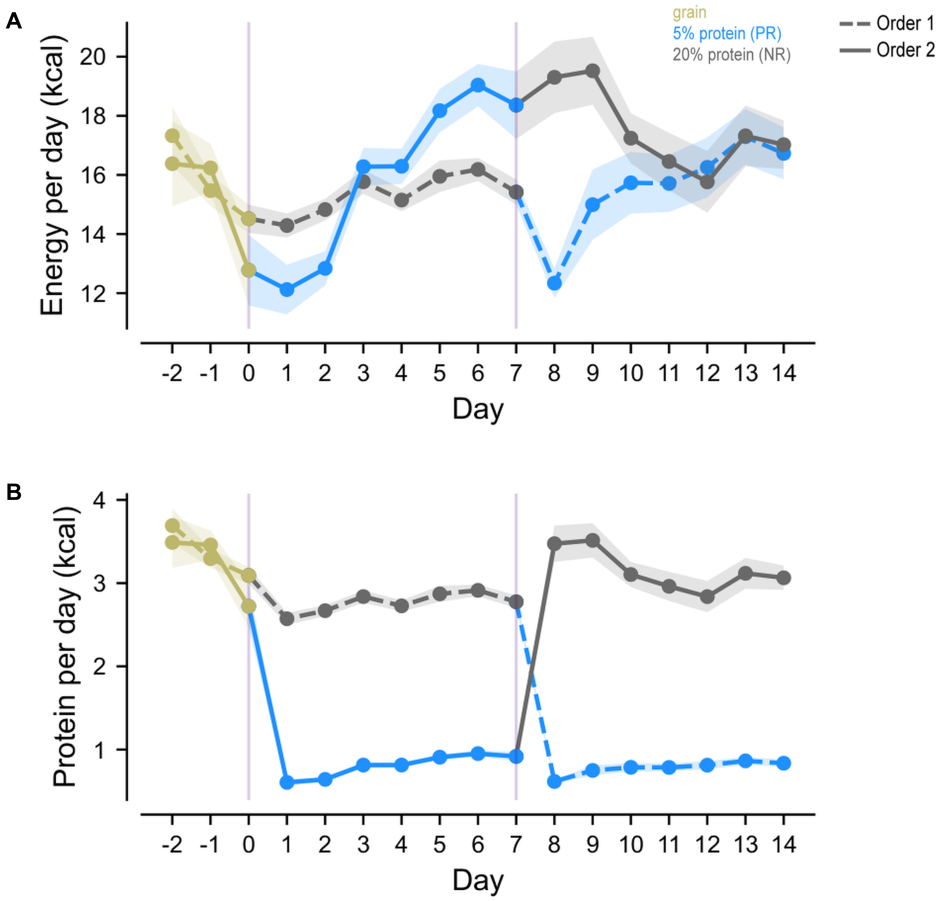


Fig. S2. In Order 2 mice, an immediate suppression of pellet intake is observed following the diet transition, likely linked to the changed caloric value of the pellets. After approximately two days, pellet intake rises steadily and remains high.

### 3.4.S1. Detailed definition of feeding microstructures using interpellet intervals and cluster-continuation analysis

To define feeding microstructures from timestamped FED3 pellet events, we used a stepwise approach. First, consecutive pellet events were grouped into feeding clusters using an interpellet interval (IPI) rule. Second, the completed cluster-size distribution was examined to describe how many pellets were contained in each feeding event. Third, neighbouring cluster-size ratios and cluster-continuation probabilities were calculated to describe the structure and progression of these events.

The IPI sequence was defined as the time difference between consecutive pellet events:

$$\mathrm{IPI}_i = t_{i+1} - t_i,$$

where $t_i$ and $t_{i+1}$ are timestamps of consecutive pellet events.

We visualised the distribution of IPIs using kernel density estimation (KDE), a non-parametric method for estimating the probability density function of a random variable:

$$\hat{f}(x) = \frac{1}{nh} \sum_{i=1}^{n} K\left(\frac{x - x_i}{h}\right),$$

where $\hat{f}(x)$ is the estimated density at point $x$, $n$ is the total number of IPI observations, $h$ is the bandwidth controlling kernel width, and $K(\cdot)$ is the Gaussian kernel function. Here, $x_i$ denotes $\log_{10}(\mathrm{IPI}_i)$. The KDE curve was used to visualise the temporal structure of pellet events and supported the use of a 60 s IPI threshold for grouping consecutive pellets into the same feeding cluster.

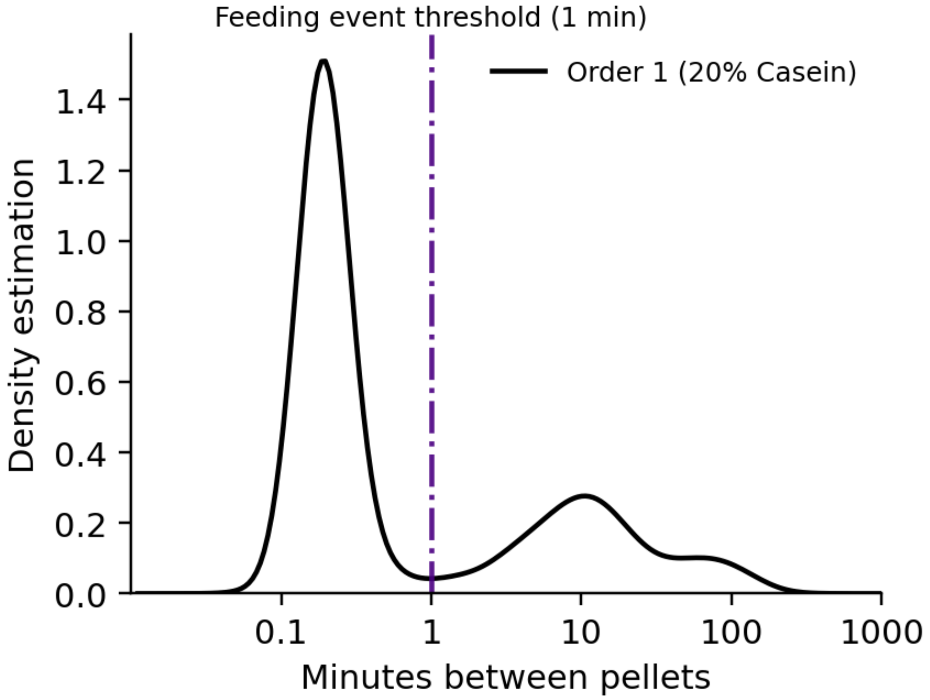


Fig. S3. KDE plot of interpellet intervals. The distribution of IPIs was used to visualise the temporal structure of pellet events and supported the use of a 60 s IPI threshold for grouping consecutive pellets into feeding clusters.

Pellets were assigned to the same feeding cluster when the interval between consecutive pellet events was no greater than 60 s. We defined:

$$\alpha_{\mathrm{IPI}} = 60 \text{ s},$$

such that if:

$$\mathrm{IPI}_i \leq \alpha_{\mathrm{IPI}},$$

then pellet $i+1$ was assigned to the same cluster as pellet $i$. If:

$$\mathrm{IPI}_i > \alpha_{\mathrm{IPI}},$$

then the preceding cluster was terminated and a new cluster was started.

After this temporal clustering step, each completed feeding cluster had a final pellet count, referred to here as cluster size. For each mouse, we counted the number of completed clusters of each size $x$. The mean frequency per mouse was calculated as:

$$y(x) = \frac{1}{N}\sum_{i=1}^{N} n_i(x),$$

where $n_i(x)$ is the number of completed clusters of size $x$ for mouse $i$, and $N$ is the number of mice.

For Order 1 mice in the NR phase ($N = 11$), the completed cluster-size distribution showed three features. First, single-pellet clusters were common, indicating many feeding events that did not continue beyond the first pellet within the 60 s window. Second, clusters containing 2–5 pellets formed the main multi-pellet range: 2- and 3-pellet clusters occurred at almost identical frequencies, and 4- and 5-pellet clusters remained common. Third, clusters larger than 5 pellets were less frequent and formed an extended-cluster range.

Table. S2. Completed cluster-size frequencies after applying the 60 s IPI threshold in Order 1 NR mice. Pooled counts show the total number of completed clusters across mice. Mean, SD and SEM are calculated across mice. Cluster sizes 1–10 are shown for readability.

| Cluster size ($x$) | Pooled count | Mean per mouse | SD | SEM |
|---|---|---|---|---|
| 1 | 1226 | 111.45 | 87.20 | 26.29 |
| 2 | 749 | 68.09 | 50.93 | 15.36 |
| 3 | 748 | 68.00 | 31.29 | 9.43 |
| 4 | 590 | 53.64 | 14.64 | 4.41 |
| 5 | 437 | 39.73 | 15.88 | 4.79 |
| 6 | 254 | 23.09 | 7.85 | 2.37 |
| 7 | 184 | 16.73 | 7.71 | 2.32 |
| 8 | 119 | 10.82 | 6.13 | 1.85 |
| 9 | 66 | 6.00 | 5.00 | 1.51 |
| 10 | 38 | 3.45 | 2.11 | 0.64 |

To summarise the overall decline in completed cluster frequency with increasing cluster size, we fitted an exponential decay model:

$$y(x) = ae^{-bx},$$

where $y(x)$ is the mean frequency of completed clusters of size $x$, $a$ is the fitted scale parameter, and $b$ is the decay rate. The fitted model was:

$$y = 144.92e^{-0.29x},$$

with $R^2 = 0.96$. This indicates that larger completed feeding clusters became progressively less frequent. The exponential decay model was used descriptively to summarise the cluster-size distribution.

To further describe the shape of the completed cluster-size distribution, we calculated neighbouring completed cluster-size ratios:

$$R_{x+1:x} = \frac{n_{x+1}}{n_x},$$

where $n_x$ is the number of completed clusters containing $x$ pellets, and $n_{x+1}$ is the number of completed clusters containing $x+1$ pellets. This ratio describes how frequent completed clusters of size $x+1$ were relative to completed clusters of size $x$.

Table. S3. Neighbouring completed cluster-size ratios after applying the 60 s IPI threshold. Ratios describe the relative frequency of neighbouring completed cluster sizes.

| **Completed cluster sizes compared** | **Calculation** | **Ratio** |
|---|---|---|
| 2-pellet relative to 1-pellet | $n_2/n_1$ | 0.611 |
| 3-pellet relative to 2-pellet | $n_3/n_2$ | 0.999 |
| 4-pellet relative to 3-pellet | $n_4/n_3$ | 0.789 |
| 5-pellet relative to 4-pellet | $n_5/n_4$ | 0.741 |
| 6-pellet relative to 5-pellet | $n_6/n_5$ | 0.581 |
| 7-pellet relative to 6-pellet | $n_7/n_6$ | 0.724 |
| 8-pellet relative to 7-pellet | $n_8/n_7$ | 0.647 |
| 9-pellet relative to 8-pellet | $n_9/n_8$ | 0.555 |
| 10-pellet relative to 9-pellet | $n_{10}/n_9$ | 0.576 |

The neighbouring cluster-size ratios showed that 2- and 3-pellet clusters occurred at nearly identical frequencies, while cluster frequencies declined across larger cluster sizes. Together with the raw cluster-size frequencies, these ratios supported treating 2–5 pellet clusters as the principal multi-pellet range and clusters larger than 5 pellets as a less frequent extended-cluster range.

We then calculated cluster-continuation probability to describe progression within ongoing feeding events. This analysis used cumulative cluster counts after applying the 60 s IPI rule. For a given pellet number $x$, the continuation probability was defined as:

$$P(\text{reach } x+1 \mid \text{reached } x) = \frac{N_{\geq x+1}}{N_{\geq x}},$$

where $N_{\geq x}$ is the number of completed clusters that reached at least $x$ pellets, and $N_{\geq x+1}$ is the number of completed clusters that reached at least $x+1$ pellets.

This cumulative calculation asks: among all feeding events that reached pellet $x$, what proportion also reached pellet $x+1$? It therefore describes the progression of ongoing events. This differs from the completed cluster-size frequency analysis, which describes how many events ended with a final size of $x$ pellets.

For each transition, continuation probabilities were first calculated separately for each mouse and summarised as mean ± SEM. We also calculated pooled continuation probabilities across all clusters to provide the total observed transition rate. For these pooled probabilities, 95% confidence intervals were calculated using the Wilson score interval:

$$\mathrm{CI} = \frac{\hat{p} + \frac{z^2}{2n} \pm z\sqrt{\frac{\hat{p}(1-\hat{p})}{n} + \frac{z^2}{4n^2}}}{1 + \frac{z^2}{n}},$$

where $\hat{p}$ is the pooled continuation probability, $n = N_{\geq x}$, and $z = 1.96$ for a two-sided 95% confidence interval.

Table. S4. cluster-continuation probabilities in Order 1 NR mice. Mouse-level values are shown as mean ± SEM. Pooled probabilities were calculated from all completed clusters across mice, with Wilson 95% confidence intervals.

| **Transition** | **Mouse-level probability** | **Pooled calculation** | **Pooled probability** | **Wilson 95% CI** |
|---|---|---|---|---|
| Reached 1 → reached 2 | $0.757 \pm 0.037$ | 3224/4450 | 0.724 | 0.711–0.737 |
| Reached 2 → reached 3 | $0.781 \pm 0.042$ | 2475/3224 | 0.768 | 0.753–0.782 |
| Reached 3 → reached 4 | $0.693 \pm 0.044$ | 1727/2475 | 0.698 | 0.679–0.716 |
| Reached 4 → reached 5 | $0.646 \pm 0.025$ | 1137/1727 | 0.658 | 0.636–0.680 |
| Reached 5 → reached 6 | $0.611 \pm 0.031$ | 700/1137 | 0.616 | 0.587–0.643 |
| Reached 6 → reached 7 | $0.618 \pm 0.029$ | 446/700 | 0.637 | 0.601–0.672 |
| Reached 7 → reached 8 | $0.556 \pm 0.044$ | 262/446 | 0.587 | 0.541–0.632 |
| Reached 8 → reached 9 | $0.535 \pm 0.029$ | 143/262 | 0.546 | 0.485–0.605 |
| Reached 9 → reached 10 | $0.605 \pm 0.050$ | 77/143 | 0.538 | 0.457–0.618 |

The continuation analysis showed that a high proportion of initiated feeding events progressed beyond the first pellet. At the mouse level, the probability of reaching pellet 2 after reaching pellet 1 was $0.757 \pm 0.037$, and the probability of reaching pellet 3 after reaching pellet 2 was $0.781 \pm 0.042$. Continuation probability then declined gradually across larger cluster sizes.

This pattern should be interpreted as stepwise cluster progression rather than as evidence that most 2-pellet events continued all the way to 5 pellets. For example, using pooled counts, the probability that a feeding event which had reached pellet 2 also reached pellet 5 was:

$$\frac{N_{\geq 5}}{N_{\geq 2}} = \frac{1137}{3224} = 0.353.$$

Thus, the continuation analysis indicates substantial stepwise continuation through the 2–5 pellet range, while the completed cluster-size distribution shows that events larger than 5 pellets were less frequent.

### Overall rationale for feeding-event classification

Overall, the classification of feeding events was based on three linked observations. First, the KDE analysis of IPIs supported the use of a 60 s temporal threshold for grouping consecutive pellet events into the same feeding cluster. Second, after applying the 60 s rule, the completed cluster-size distribution separated isolated 1-pellet events from multi-pellet events. Third, the 2–5 pellet range accounted for the majority of multi-pellet events, while events larger than 5 pellets formed a less frequent extended-cluster range.

The number of completed 2–5 pellet events was:

$$749 + 748 + 590 + 437 = 2524.$$

The total number of multi-pellet events was:

$$N_{\geq 2} = 3224.$$

Thus, 2–5 pellet events accounted for:

$$\frac{2524}{3224} = 0.783,$$

or approximately 78% of all multi-pellet feeding events. Events larger than 5 pellets accounted for:

$$\frac{700}{3224} = 0.217,$$

or approximately 22% of all multi-pellet feeding events.

Based on these observations, the final operational classification used for downstream analysis was:

$$\begin{cases} 1 \text{ pellet} & = \text{snack}, \\ 2\text{–}5 \text{ pellets} & = \text{meal}, \\ > 5 \text{ pellets} & = \text{feast}. \end{cases}$$

Table. S5. Feeding-event classes after applying the 60 s IPI threshold. Values show the number of completed clusters in Order 1 NR mice. The feast category includes all clusters containing more than 5 pellets.

| **Class** | **Definition** | **Pooled count** | **Mean per mouse** | **SEM** |
|---|---|---|---|---|
| Snack | 1 pellet | 1226 | 111.45 | 26.29 |
| Meal | 2–5 pellets | 2524 | 229.45 | 20.79 |
| Feast | > 5 pellets | 700 | 63.64 | 7.92 |

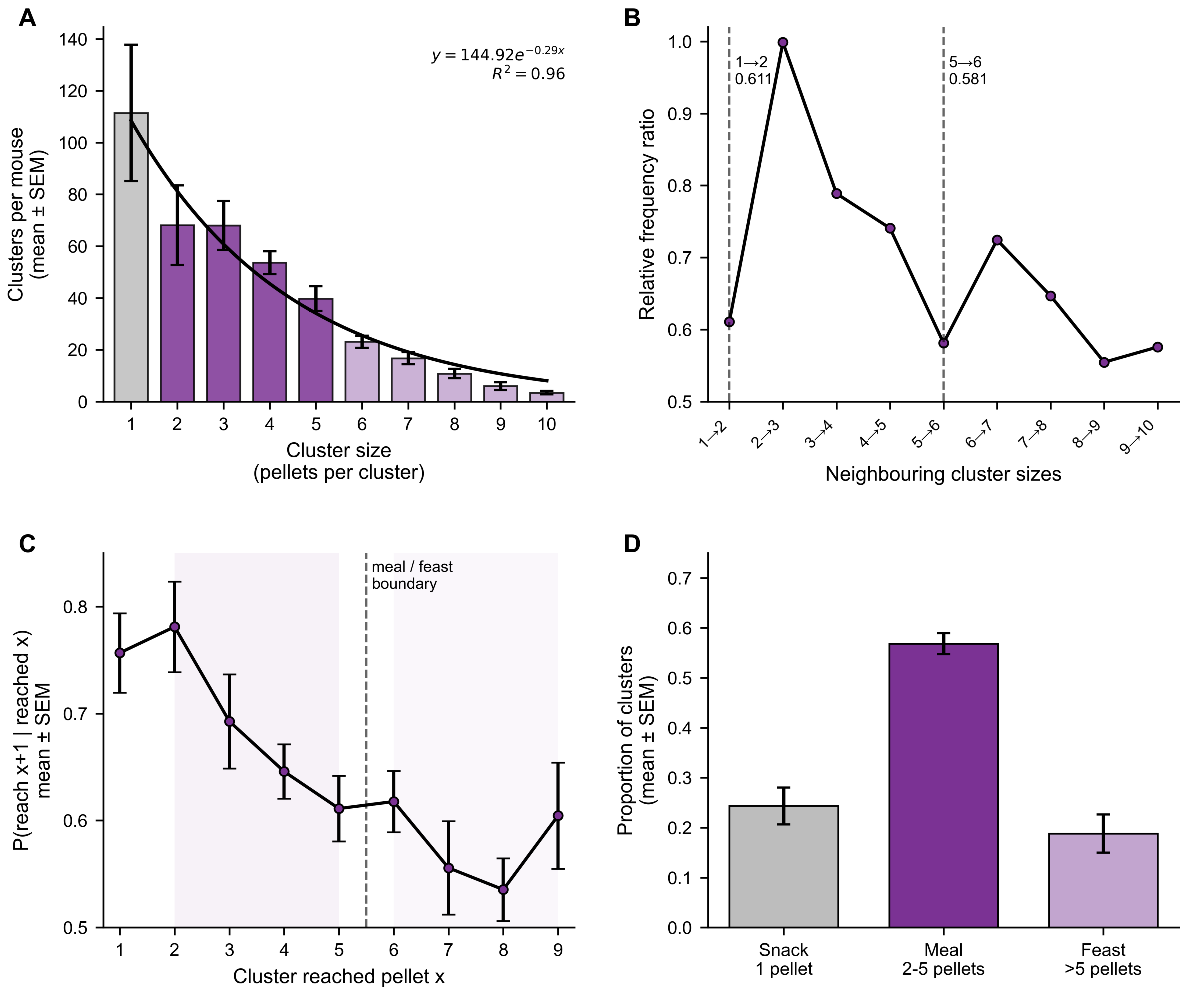


Fig. S4. Feeding-event cluster sizes and cluster-continuation probability. (A) Completed cluster-size frequencies after applying the 60 s interpellet interval threshold, shown as mean cluster counts per mouse with an exponential decay fit. (B) Operational grouping of completed clusters into snacks, meals and feasts. (C) cluster-continuation probability, calculated as the proportion of clusters reaching at least $x$ pellets that also reached at least $x+1$ pellets. The dashed line marks the operational boundary between meals and feasts.

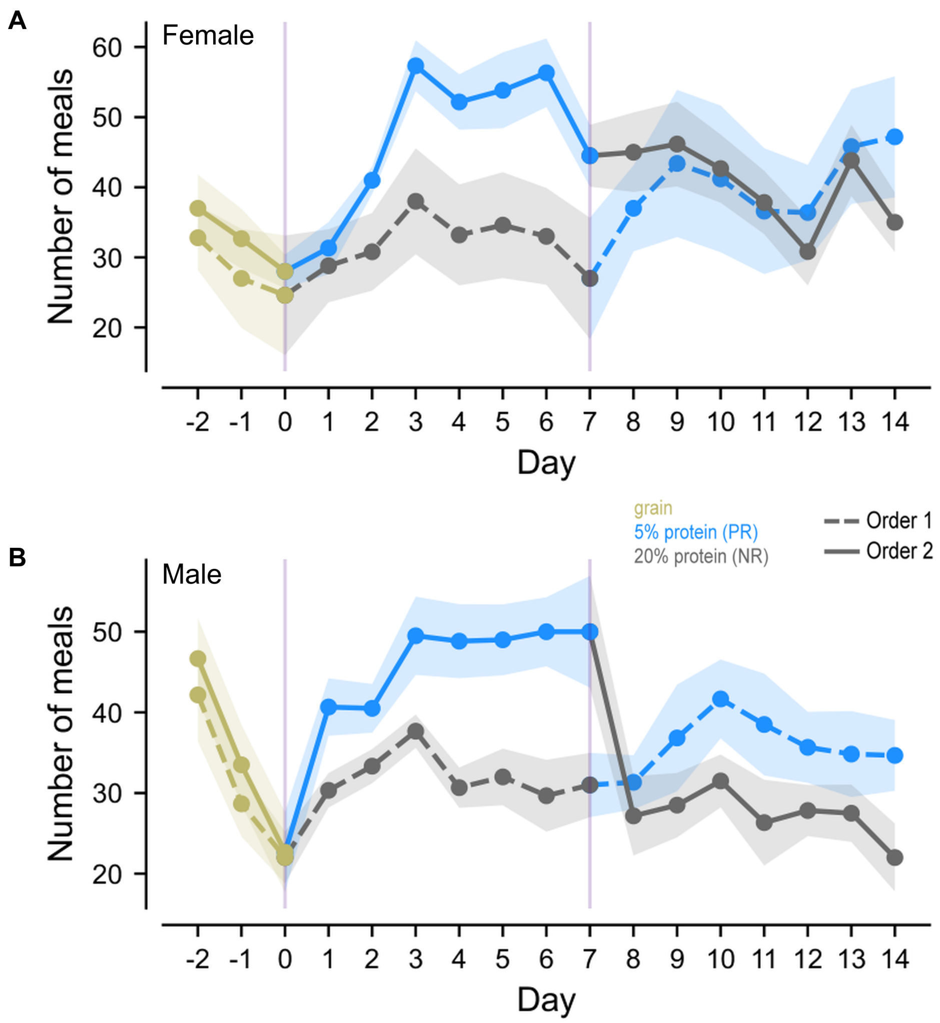


Fig. S5. Day-to-day number of meals (2–5 pellets) shown separately for female (A) and male (B) mice. Data are presented as mean ± SEM. Dashed lines indicate Order 1 and solid lines indicate Order 2. Colours denote diet (grain, 5% protein (PR), and 20% protein (NR)). These sex-specific plots correspond to the pooled analysis shown in Fig. 5B.

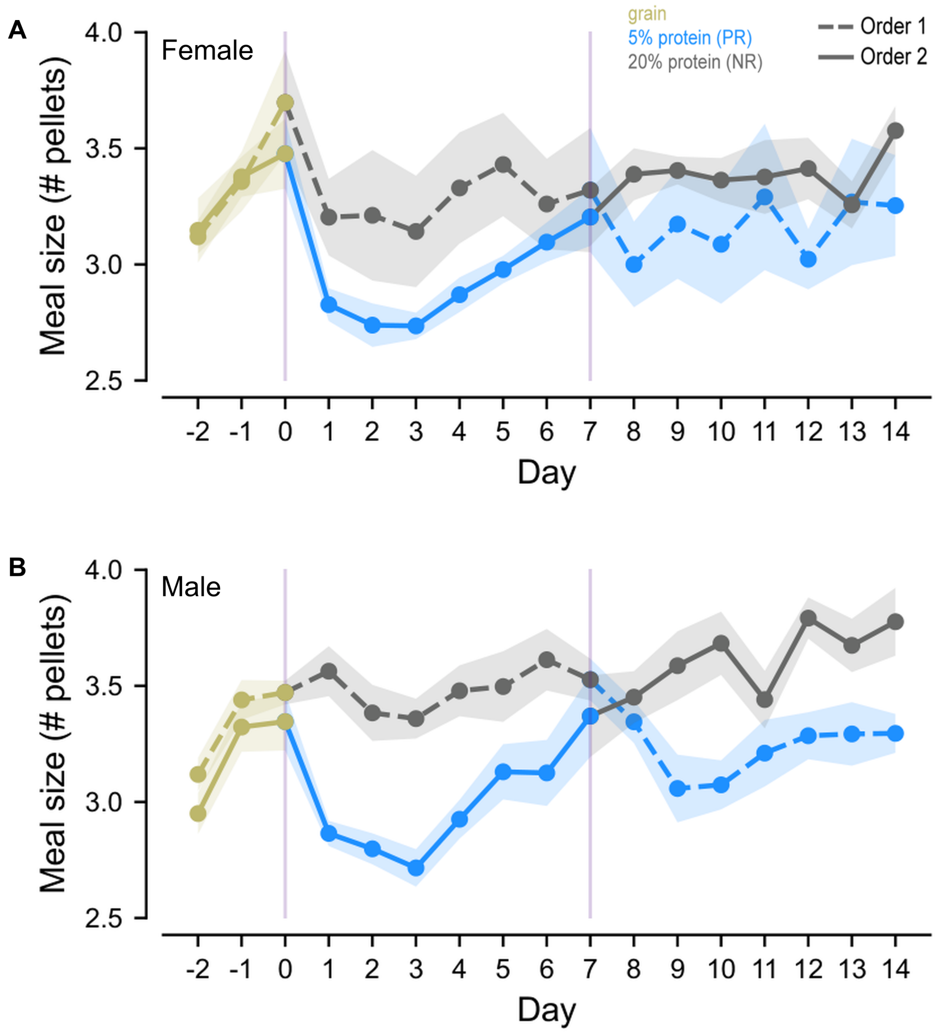


Fig. S6. Day-to-day meal size (pellets per meal) shown separately for female (A) and male (B) mice. Data are presented as mean ± SEM. Dashed lines indicate Order 1 and solid lines indicate Order 2. Colours denote diet (grain, 5% protein (PR), and 20% protein (NR)). These sex-specific plots correspond to the pooled analysis shown in Fig. 5D.

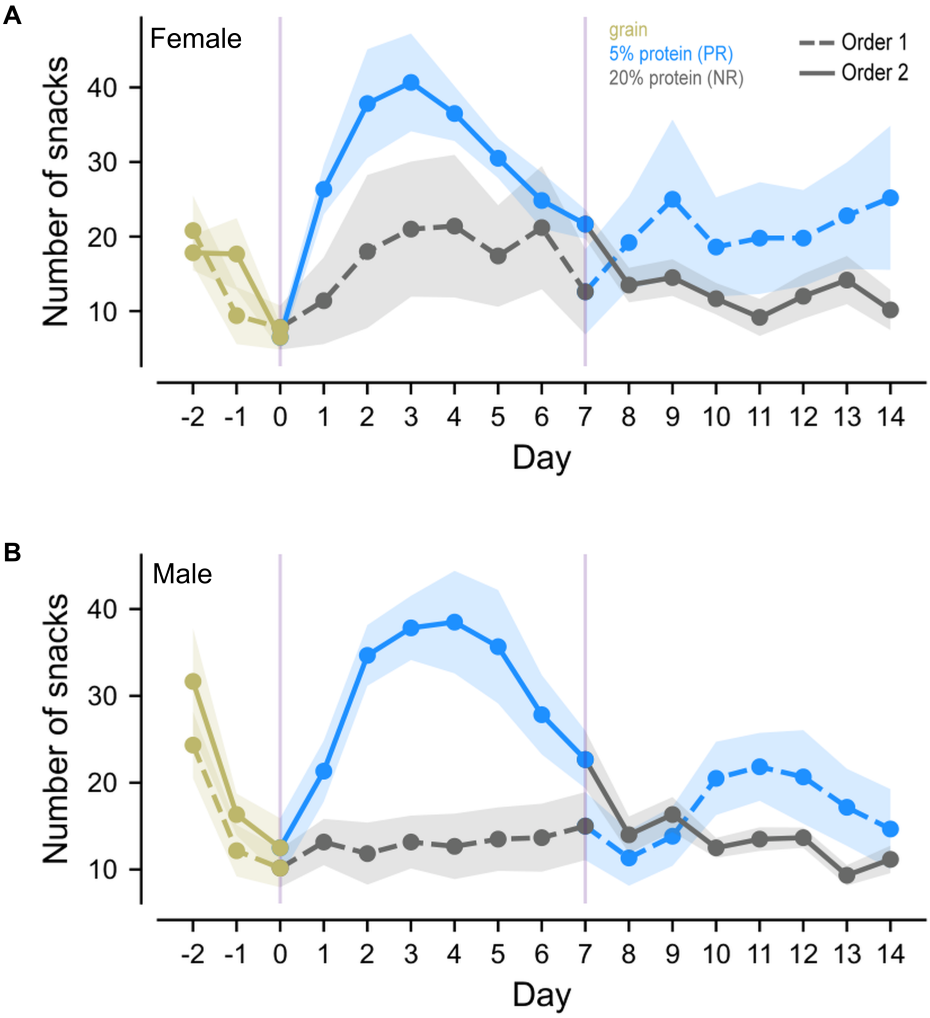


Fig. S7. Day-to-day number of snacks shown separately for female (A) and male (B) mice. Snacks were defined as eating events consisting of one pellet. Data are presented as mean ± SEM. Dashed lines indicate Order 1 and solid lines indicate Order 2. Colours denote diet (grain, 5% protein (PR), and 20% protein (NR)). These sex-specific plots correspond to the pooled analysis shown in Fig. 6B.

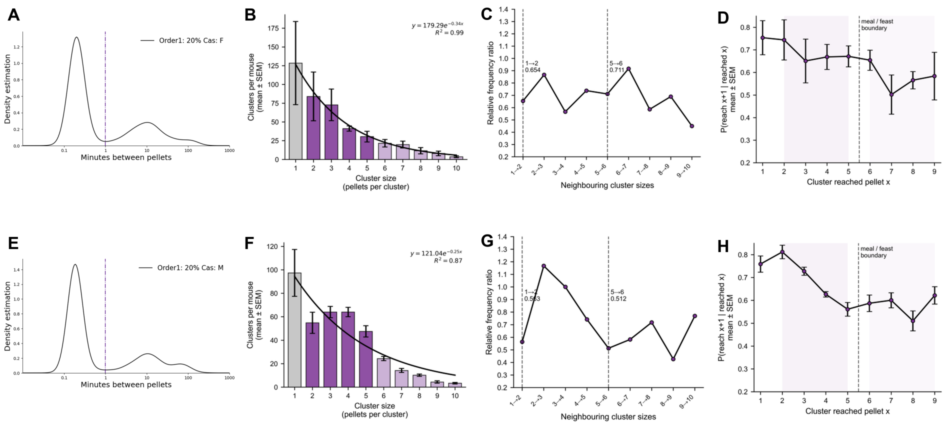


Fig. S8. Sex-stratified feeding-event structure in Order 1 mice during the NR reference phase. Female (A–D; $n = 5$) and male (E–H; $n = 6$) data are shown separately. (A, E) Inter-pellet interval distributions; dashed lines indicate the 60 s event threshold. (B, F) Completed cluster-size frequencies (mean ± SEM) with exponential fits. (C, G) Ratios of neighbouring completed cluster sizes ($n_{x+1}/n_x$). (D, H) Cluster-continuation probabilities ($P(\text{reach } x+1 \mid \text{reached } x)$), shown as mean ± SEM. Dashed lines and shading indicate the common operational categories derived from the pooled reference data: snacks (1 pellet), meals (2–5 pellets), and feasts (> 5 pellets).
